# Manipulating Environmental Affordances via Intrinsic Measurement: Evidence from Sensorimotor Brain Dynamics

**DOI:** 10.1101/2025.01.05.631377

**Authors:** Sheng Wang

## Abstract

According to the view of ecological psychology, affordances are perceived directly through the interaction between the observer and the environment, and thus body-scaled. Considering the actions from different object-agent distances, Tosoni and colleagues have provided a conceptual distinction concerning two types of affordances: macro affordances (also called environmental affordance) and micro affordances. Both of these two types of affordances have been investigated in psychological research since the beginning of this century. However, most of the environmental affordances have not yet been manipulated with a body-scaled setting in the prior research protocol. Inspired by the intrinsic measurement introduced by ecological psychologist Warren (1984) and focusing on environmental affordances, the present study adapted the task of Djebbara et al. (2019) by defining transitional affordance as the body-door height ratios. The current results from subjective data, EEG data and behavioral data showed that affordance can be manipulated intrinsically and covaried with agents’ experience, perception and motor interaction. The perception of environmental affordances was based on intrinsic optical information for the relationship between environmental properties and capacities of the observer’s own action system. It thus provides empirical support for the need to link ecological psychology and neuro-architecture in experimental protocols.

## **1.** Introduction

### 1.1. Background and literature review

The concept of affordances, introduced by ecological psychologist J. J. Gibson in the late 1960s and 1970s, refers to the opportunities for action offered by the environment to an agent (Gibson, 1966, 2014). According to our environmental experience, a leaf provides shelter for a wasp but not large enough for a dog; a chair with 50 cm height is sit-able for an adult but not for a baby. Here, the shelter-ability of the leaf is an affordance for the wasp; and the sit-ability of the chair is an affordance for the adult. Thus, affordances have been traditionally understood as relational or agent-relative properties: affordances are “relations between the abilities of organisms and features of the environment” [(Chemero, 2003), p. 189; see also (Chemero, 2013)]. Consistent with this principle, in a landmark study Warren quantified affordance in an intrinsic manner to support the relational understanding of affordances. Warren (1984) found that the boundary between climbable and unclimbable stairways corresponds to a fixed ratio between riser height and leg length. That is, instead of the stairway having the affordance of “climbability” on its own, the affordance is rather a relational property—one to which participants in Warren’s study were found to be perceptually sensitive. This research introduced a methodology called intrinsic measurement to quantify affordances, as the unit of climbability is not an extrinsic measure, such as centimeters, but rather an intrinsic one tied to the body-environment relation, dependent on leg lengths. (Warren, 1984).

Considering the actions from different object-agent distances, Di Marco et al. (2019) raised a term “macro-affordance” to reflect the parallelism with the well-known term “micro-affordance”. Macro affordances or called as environmental affordances were used interchangeably in their work, reflecting the impact of affordance on locomotion in the coverage of extrapersonal distance (Altomare et al., 2021; Bonner & Epstein, 2017; Di Marco et al., 2019; Djebbara et al., 2019; Tosoni et al., 2021, 2023); whereas the micro affordances describe the impact of affordance on action within the hand-related reachable/peripersonal space (Costantini et al., 2010; Ellis & Tucker, 2000; Feng et al., 2024). Both micro and macro affordances have captured attention within contemporary psychological research.

In order to quantify the impact of affordance information on action, yet from a micro perspective, scholars traditionally formalized this impact as affordance-facilitation/compatibility effects. Specifically, this effect refers to the phenomenon where an agent’s motor response is optimized or enhanced when the spatial position of an object aligns with an agent’s sensorimotor capacities to interact with it, based on the perception of the object’s affordances (see Costantini et al., 2010; Derbyshire et al., 2006; Ellis & Tucker, 2000; Tucker & Ellis, 1998, 2001, 2004; for theoretical papers see Ferretti, 2016, 2021; Zipoli Caiani, 2014). To the best of the author’s knowledge, this (facilitation) effect was first termed by Ellis and Tucker in 2000 as micro-affordance (Ellis & Tucker, 2000). Yet their landmark study was conducted 2 years ago. Using a stimulus–response compatibility (SRC) paradigm, Tucker and Ellis (1998) required participants to decide as fast as possible whether objects displayed on a computer monitor were upright or inverted, with objects presented in orientations compatible with either right or left-hand grasps. Responses were faster when the object’s handle aligned with the responding hand, even though the handle of the object was irrelevant to the judgment about the objects’ vertical orientation. This demonstrated that perceived affordances can automatically facilitate grasp-related actions, even in the absence of explicit intentions to act (Tucker & Ellis, 1998). And then Costantini et al. (2010) found the micro-affordance effect only when the object was located within the peripersonal space. Inspired by some above studies concerning micro-affordance, a latest study by Feng et al. (2024) investigated how object perception is influenced by the size of objects relative to the human body. With the evidence from behavior, brain, and large language models, they concluded that only objects within the body size range are capable of being manipulated, and this manipulability is a causal premise for objects to offer affordance in the eyes of an organism (Feng et al., 2024). Generally, although micro-affordance research took into consideration an individual’s sensorimotor capacities, even including body size as a metric for affordance interaction, they potentially neglected the agents’ active exploration in the surrounding environment and the enactive perception of environmental affordances. This motive promoted the research regarding macro/environmental affordance, with the contribution of scholars from multi-fields like neuro-architecture (Djebbara et al., 2019, 2022; S. Wang et al., 2022), environmental psychology (Di Marco et al., 2019; Tosoni et al., 2023) and spatial navigation (Bonner & Epstein, 2017; Gregorians & Spiers, 2022; Harel et al., 2022).

While sharing a similar objective with micro-affordance studies, scholars in the field of macro affordance have explored the impact of perceived affordances on action, but from an environmental perspective. For example, based on the finding that an affordance facilitation effect for a walking-related action was also observed in the far extrapersonal/environmental space, Di Marco et al. (2019) first raised a term “macro-affordance” to reflect the parallelism with the well-known term “micro-affordance”. Tosoni et al. (2021) extended this landmark study to investigate the neurophysiological mechanism of the "macro-affordance" effect using an incidental Go/NoGo priming paradigm. Participants responded to repeated presentations of environmental pictures by executing a walking-related action (a footstep) for Go trials, while refraining on NoGo trials when the prime and target pictures differed. The study revealed that distant (vs. near) objects in extrapersonal space facilitated footstep actions, demonstrating a macro-affordance effect. And the source localization of this effect was observed in sensory-motor regions of the dorso-medial fronto-parietal network (Tosoni et al., 2021). Focusing on transitional affordances in a built environment, previous work by our team has shown that cortical dynamics covary with perceiving and transitioning differently wide doors (Djebbara et al., 2019) according to different sensorimotor stages, indicating that the perceptual processing of environmental affordances is flexible and time-varying (S. Wang et al., 2024). Taken together, during agents’ active exploration in the surrounding environment, the affordances perception can be extended over different sensorimotor stages (for review papers see Djebbara et al., 2022; Pezzulo & Cisek, 2016). Although the research regarding macro/environmental affordance reflected the sensorimotor dynamics of affordance perception in the surrounding environment, most of the experimental protocol settings were not intrinsic or body-scaled.

### 1.2. Research question and hypothesis

Taken together, affordance impacts action not only on the level of object manipulation but also on the level of space exploration. The affordances involving object manipulation invite actions within the body size range, whereas affordances can also emerge through active exploration in the surrounding environment, referred to as micro-affordances and macro-affordances, respectively. However, due to different theoretical backgrounds and experimental paradigms, previous experimental protocols remains us an impression of dualistic antagonism or conceptual tension: macro-affordances are situated at an extrapersonal distance, making them difficult to define within a body-scaled setting, while intrinsic affordances rarely extend into the surrounding environment, thereby neglecting the enactive perception and the sensorimotor dynamics involved as agents explore the environment. Therefore, the present study investigated whether a type of affordance exists that combines properties of both the body scale and the macro level. It is hypothesized that environmental affordances can be manipulated through an intrinsic setting, and that varying levels of affordance manipulation may prime agents’ experience, motor interaction, and perception. If this were the case, then affordances should systematically modulate subjective data, behavior and brain activity. Using mobile brain/body imaging (MoBI) approach (Jungnickel et al., 2019; Makeig et al., 2009), the current research aimed to provide a paradigm extension of the operational definition of affordance, which goes beyond the notion of both micro-intrinsic affordance and macro-extrinsic affordance, by offering the one of ‘macro-intrinsic affordance’. Focusing on this research target, in order to operationally define environmental affordance in an intrinsic manner, the present study adapted the task of Djebbara et al. (2019) by defining transitional affordance as the body-door height ratios. To this end, participants actively walked through a 3D virtual environment with the task to pass differently high doors to go from one room to an adjacent room. In order to investigate the sensorimotor brain dynamics of environmental affordances, during the transition display, early visual evoked potentials and motor related potentials were collected as components to correspond visual features extraction of affordances and later sensorimotor coupling processing respectively. Besides EEG data, subjective rating and behavioral data were also collected.

## 2. Method

### 2.1. Participants

Twenty participants (eleven females) without history of neurological pathologies were recruited from social media and a participant pool of the Technical University of Berlin, Germany. Their mean age was 26.58 years (σ = 5.409). Their mean body height was 170.79 centimeters (σ = 4.685). All participants signed the consent form, which was approved by the Ethics Committee of the Technical University of Berlin. After each data collection, the participants received either monetary compensation (12 €/hour) or accredited course hours. All participants had normal or corrected-to-normal vision, and none had a specific background in architecture. One male participant was excluded from the study due to technical issues.

### 2.2. Experimental paradigm and procedure

The experiment was conducted at Berlin Mobile Brain/ Body Imaging Laboratories (BeMoBIL) in an experimental room with an area of 160 m^2^. The size of the virtual space was 11.5 m × 5 m, with room sizes of 6.5 m × 5 m for the first room and 4.85 m × 5 m for the second room and 0.15 m × 5 m for the wall between them. These two adjacent rooms were connected through a door. Wearing a head mounted VR, Participants’ task was to physically walk through this virtual transition. Participants wore a backpack, which held a high-performance gaming computer (Zotac ZBOX Magnus EN374070C) to render the VR environment, along with an EEG amplifier system (ANT Neuro, eego™sports). The VR experience was delivered using a HTC Vive Pro headset (1440 × 1600 per-eye resolution, 90 Hz refresh rate, 98° vertical and horizontal field of view, 800 g weight with headstrap) connected to the Zotac computer. Participants used a HTC Vive controller to interact with the virtual environment, which was built using Unity (Figure 1).

**FIGURE 1.**
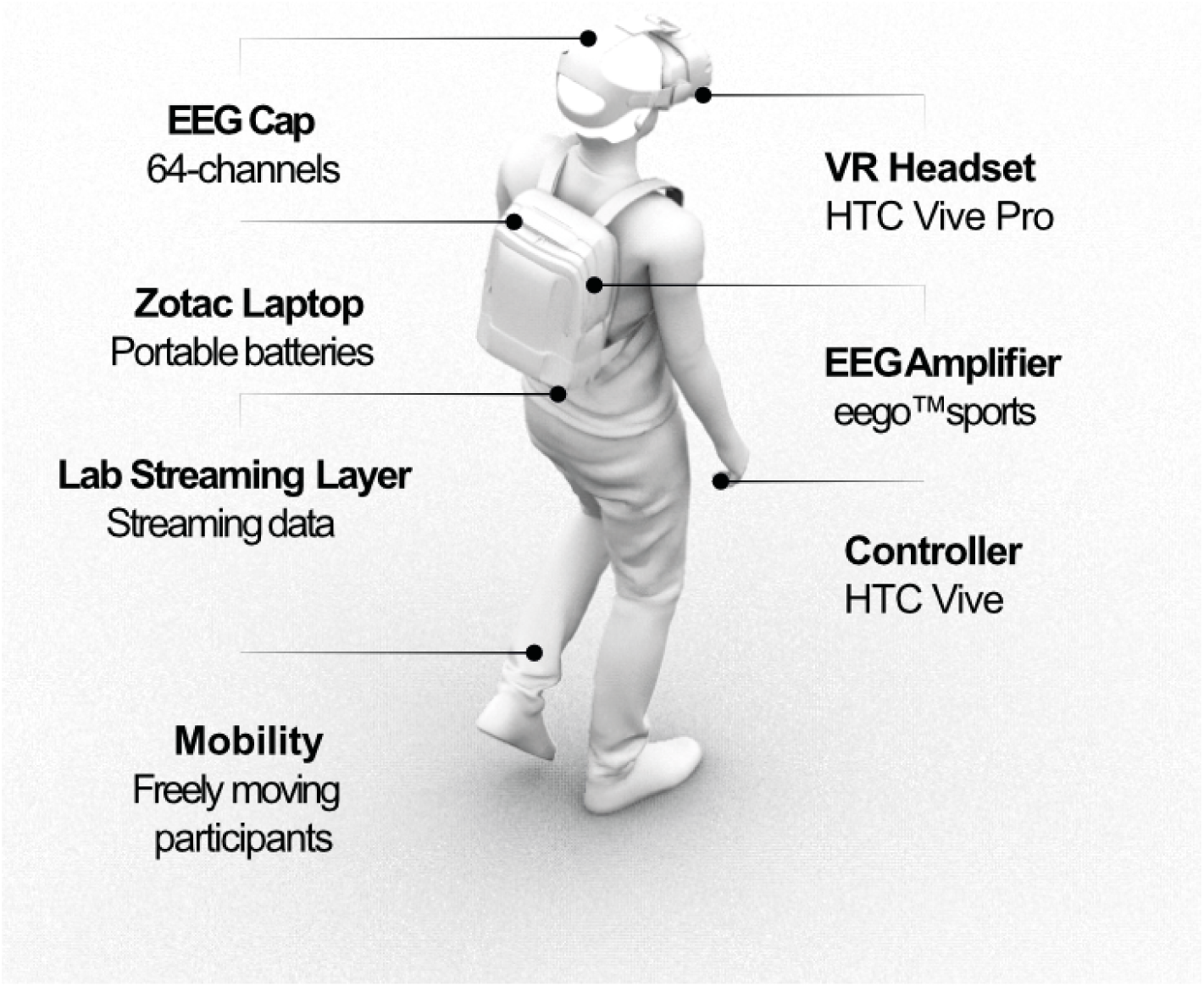
MoBI Setup. Participants carried a high-performance gaming computer (Zotac) in a backpack, powered by two batteries. An EEG amplifier (ANT eegoSports) was also attached to the backpack and connected to the Zotac computer. They wore a 64-channel cap and also a HTC Vive Pro headset. This configuration enabled free movement during data recording.

To operationally define environmental affordance intrinsically, the present paradigm adapted the task of Djebbara et al. (2019) by defining transitional affordance as body-door height ratios. The doors varied in height from high (120% body height, easily passable) to mid (95%, passable) to low (60%, difficult to pass), presented in a pseudorandomized order. The height variations manipulated transition affordance, with low and mid-height doors affording opportunities for body posture adjustments. Participants performed a forewarned (S1-S2) paradigm with two movement instructions (pseudorandomized 50/50) to either walk through the door (‘Go’ trial) or not (‘NoGo’ trial). In Go trials, the participants were required to walk from the first room to the second room and to pick up a token, whereas in the other half of the trials (NoGo), they simply judged the experience of the first room they were located in without moving to the second room. As mentioned in our previous paper (S. Wang et al., 2024), the current experiment does not follow the traditional Go–NoGo task paradigm in which subjects should respond fast to frequently presented ‘Go’ targets while ignoring rare ‘NoGo’ stimuli (Aron & Poldrack, 2005). Instead, the terms ’Go trials’ and ’NoGo trials’ were used to refer to the imperative stimulus in the respective trials, following the protocol established by Djebbara et al. (2019).

The experiment was a 2 × 3 repeated-measures design with factors of imperative stimulus (Go, NoGo) and door height (low, mid, high). Each participant completed 240 trials, with 40 trials per condition. In each trial, participants started in a dark environment on a predefined square (Figure 2). After a random interval (mean = 5 s, SD = 2 s), the ‘lights’ turned on, revealing a room with a closed door. Following another random delay (mean = 4 s, σ = 2 s), the door color changed to green (Go trial) or red (NoGo trial). In Go trials, participants were instructed to walk toward the door, which slid open upon reaching it. They then passed through, found a red rotating ring in the second room, and touched it to earn a 1-cent bonus. After returning to the first room, they provided a subjective rating using the self-assessment manikin (SAM) questionnaire. However, in the case of a red door (NoGo), participants stayed in the first room and provided a subjective rating without passing through the door. After the rating, the participants pressed the trigger button on the controller to start the next trial.

**FIGURE 2.**
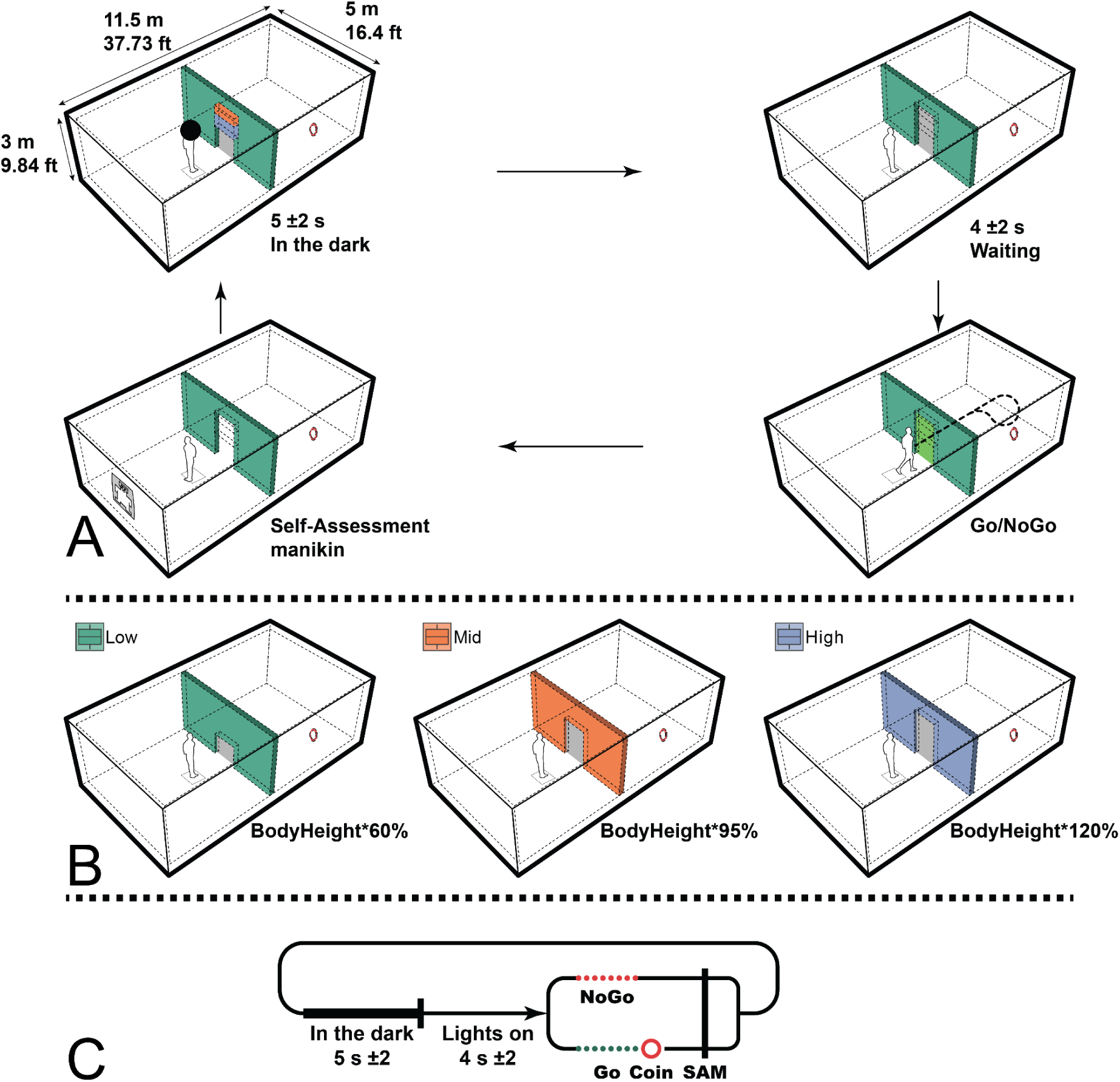
(A) Participants were instructed to view the VR screen. First, in the start square within the Unity environment, the screen was pure black for 5 s (σ = 2 s). Second, the moment the light was on, the black sphere disappeared and the participants perceived the door they had to pass. The door height was either low (body height × 60%), mid (body height × 95%) or high (body height × 120%). All doors in the experiment had the same colour and only varied with respect to the distance of the horizontal lines indicating the door height. Door colours in the figure were not presented to the participants and serve only as illustration purposes. Then they waited for the imperative stimulus, either a green door (go) or a red door (NoGo), for 4 s (σ = 2 s). In the case of Go, the participants were instructed to walk toward the door. Upon entering the second room, the participants had to find and virtually touch a red rotating coin. Successfully touching the coin earned participants an additional 1 cent as a bonus. The bonus number was displayed on the small virtual screen of the controller and updated regularly. The participants then returned to complete the virtual SAM questionnaire. In the case of NoGo, the participants were instructed to turn around and complete the virtual SAM questionnaire. Finally, after completing the SAM questionnaire, the participants were instructed to press the trigger button on the controller to restart the next trial. **(B)** There were three dimensions of door height conditions: low, body height × 60% (green); mid, body height × 95% (red); high, body height × 120% (purple). Note the colour code for each door as used throughout the paper. **(C)** A diagrammatic timeline depicting the sequence of events for a single trial.

In each Go trial, participants were instructed to move through the door into the second room, regardless of its passability, using only their legs and without arm support to ensure motor consistency across trials. If participants touched the surrounding walls, they were warned with a red color and auditory signal, indicating a failure to pass and the loss of the bonus (the red rotating ring in the second room disappeared). Participants first completed a training session to familiarize themselves with the environment and conditions. The experimenter communicated with participants via a camera and microphone from a control room, with five breaks in total: two mandatory (at least 2 minutes) and three optional.

### 2.3. Self-assessment manikin

To assess participants’ subjective experience, they completed a virtual SAM questionnaire after each trial. The SAM is a pictorial assessment of valence, arousal and dominance on a 5-point Likert scale (Bradley & Lang, 1994). Participants self-assessed their state by pressing a trigger button corresponding to a number on the scale; the first press allowed for changes, and a second press confirmed the selection. The SAM data were analyzed using a 2 × 3 factorial repeated-measures ANOVA with the factors imperative stimulus (Go and NoGo) and door height (low, mid and high). Post hoc analyses were computed using the least significant difference (LSD) test to correct for pairwise comparisons.

### 2.4. Door approaching time

In order to investigate whether the willingness to pass the door differs according to different door heights, experimenters obtained the door approaching time for all Go trials. The time when participants began moving out of the starting area and the onset of the doors sliding open were recorded, allowing for the calculation of the approach time for each Go trial.

### 2.5. EEG recording and data analysis

All data streams, including experimental events such as participants’ movement and location in the Unity environment, stimulus markers and button pressing, SAM answers and EEG data, were recorded and synchronised using Lab Streaming Layer (LSL; (Kothe et al., 2014)).

EEG data were recorded with a 64-channel EEG system (ANT Neuro, eego™sports), sampled at 500 Hz and referenced to CPz. The amplifier was a true DC amplifier and no analog or digital filters were applied to the raw data. The ground electrode was placed at AFz (anterior frontal midline) according to the 10-10 system (Oostenveld & Praamstra, 2001). Experimenters used wet electrodes and impedances were kept below 10 kΩ. Data analysis was conducted using MATLAB (MathWorks), the EEGLAB toolbox (Delorme & Makeig, 2004) and custom scripts. Before data preprocessing, all training and resting data sessions were excluded. All preprocessing steps were based on the BeMoBIL pipeline (Klug et al., 2022), first down-sampling the data to 250 Hz and then applying the ZapLine-plus function to remove spectral peaks at 50 Hz, corresponding to the power line frequency (Klug & Kloosterman, 2022). Next, bad channels were detected and interpolated in a repeated process using the clean raw data plugin of EEGLAB. To ensure a reproducible bad channel detection, the researcher ran the clean_artifacts function, with a recommended number of 10 iterations. During each iteration, channels with a correlation to a random sample of other channels were marked for removal in case the correlation was below 0.8 for more than 50% of the processed data. And subsequently bad channels were interpolated using a spherical spline interpolation. As a final step, the data were re-referenced to the average of all scalp channels, excluding the EOG channel. Subsequently, on each cleaned dataset, we computed an independent component analysis (ICA) using an adaptive mixture independent component analysis (AMICA) algorithm (Palmer et al., 2011) with the recommended parameter values from Klug and Gramann (2021). Prior to AMICA, the researcher performed high-pass filtering with a 1.5 Hz cutoff frequency as recommended by Klug and Gramann (2021) for mobile EEG data. Then the AMICA decomposition was computed using one model and 10 iterations of data rejection with a rejection threshold of 2.7 standard deviations from the log-likelihood of the explained data during the decomposition. Afterwards, an equivalent current dipole model was computed using the DIPFIT toolbox of EEGLAB. Independent components (ICs) were automatically classified using ICLabel (Pion-Tonachini et al., 2019). In the current mobile case, ICs were labeled using the ICLabel ‘lite’ classifier (for the methodological background see Klug & Gramann, 2021). Subsequently, the researcher removed ICs reflecting eye-movement activity (mean = 3.42, SD = 1.17). The computed AMICA information including rejections and dipole fitting was copied back to the initial preprocessed, but unfiltered dataset, considering that final EEG measures (e.g., ERPs) may require a lower cutoff-filtering on the EEG data (Klug et al., 2022; Klug & Gramann, 2021). As a very last step, the data is cleaned with the ICLabel classifier ‘lite’, as mentioned above.

The cleaned datasets after AMICA were band-pass filtered between 1 and 40 Hz, and two sets of epochs were extracted with the first set for the onset of the first room (lights on) and the second set with onset of the imperative stimulus (Go/NoGo). All epochs were extracted from 1000 ms before to 1000 ms after stimulus onset for low, mid and high door trials separately. And all epochs were extracted from 500 ms before to 1500 ms after stimulus onset for Go/NoGo imperatives separately. Then, all epochs were baseline corrected using a 200 and 0 ms pre-stimulus interval and cleaned using the bemobil_reject_epochs function to rank epochs with respect to their noise level and removing a fixed percentage of the worst ranking epochs (Klug et al., 2022). Each epoch was evaluated using four measures that were normalised by their median across epochs: (i) mean of channel means (weight = 1), (ii) mean of channel SDs (weight = 1), (iii) SDs of channel means (weight = 1) and (iv) the SD of channel SDs (weight = 1). Each epoch then received a final summed score, and the epochs were sorted according to that score. Finally, up to 15% of epochs were removed (mean = 35.58, SD = 0.692) accordingly.

The visual evoked potentials and motor-related cortical potentials (MRCPs) were analyzed both globally at central midline electrodes (Fz, FCz, Cz, Pz, POz, and Oz). These electrodes locations cover brain regions involved in processing visual information and regulating motor actions, however, only four channels (Fz, FCz, POz, and Oz) are discussed here, considering the prior studies on affordances (Bozzacchi et al., 2012, 2015; Djebbara et al., 2019; Goslin et al., 2012; Rowe, 2018). Although a typical P1-N1 complex was not observed at the grand average level, two pronounced peaks (P140, N260) were identified at posterior electrodes (Pz, POz, and Oz) after the ’lights on’ stimulus onset through visual inspection. The researcher also identified the N140 and P260 components at the anterior electrodes (Fz, FCz, and Cz). Individual peak latencies were determined considering both EEG polarity and a time range of 50 to +50 ms around the group-level peak latencies. Therefore, the search windows for individual peaks ranged from 90 to 190 and 210 to 310 ms for the above early ERP peaks, respectively. To ensure robust statistical analysis, each negative or positive peak was exported, computing the mean from the data covering the peak plus/minus 3 sample points before and after the peak. A global ANOVA analysis was conducted on the amplitudes with the latency of 140 ms and 260 ms respectively. The present experiment used a 3 × 6 repeated-measures ANOVA with within-subjects factors door height (low, mid and high) and electrodes. However, the present study was cautious with interpreting main effects since polarity reversals may potentially cancel each other out and thus focused on the interaction effect. Post hoc analyses were computed using the LSD test to correct for pairwise comparisons.

For the motor-related cortical potentials (MRCPs), the current study focused on the post-imperative negative variation (PINV) according to the findings of Djebbara et al. (2019). This component can be considered a postimperative negative variation reflecting the motor anticipation or the readiness to act related to onset of an imperative stimulus (Casement et al., 2008; Djebbara et al., 2019; Elbert et al., 1982). According to the visual inspection at the grand average level and classical studies (Diener, Kuehner, et al., 2009; Verleger et al., 1999), here the PINV component was calculated as the mean amplitude in the range of 1100-1400 ms after stimulus onset (imperative stimulus: Go/NoGo). The data of PINV was analysed using a 2 × 3 × 6 factorial repeated-measures ANOVA with the factors imperative stimulus (Go and NoGo), door height (low, mid, and high), and electrode location (Fz, FCz, Cz, Pz, POz, and Oz). However, the present study was cautious with interpreting main effects since polarity reversals may potentially cancel each other out and thus focused on the interaction effects. Post hoc analyses were computed using the LSD test to correct for pairwise comparisons. The present paper described the p value lower than 0.05 as a significant result and the p value higher than 0.05 as a non-significant result.

## 3. Results

To manipulate the environmental affordance in an intrinsic protocol and investigate its sensorimotor mechanisms, experimenters collected subjective, behavioral, and electrophysiological data, with a focus on electrophysiology. All data underwent statistical processing and are reported here in the order processed.

### 3.1. Subjective data: SAM ratings (figure 3)

The SAM questionnaire was completed for both Go and NoGo trials in all door height conditions. A 2 × 3 factorial repeated-measures ANOVA was used with the factor imperative stimulus (Go and NoGo) and the factor door height (low, mid and high) for each dimension of the SAM questionnaire (see Figure 3). For arousal, there was a significant main effect of the factor door height (F_2,36_ = 3.496, P = 0.041, partial η^2^ = 0.163) as well as a significant main effect of the factor imperative stimulus (F_1,18_ = 6.074, P = 0.024, partial η^2^ = 0.252). And the interaction of both factors was not significant (F_2,36_ = 0.938, P = 0.401). For valence, there was a significant main effect of the factor door height (F_2,36_ = 8.444, P < 0.001, partial η^2^ = 0.319) as well as a significant interaction of both factors (F_2,36_ = 12.477, P < 0.001, partial η^2^ = 0.409). There was no main effect of the factor imperative stimulus (F_1,18_ = 3.344, P = 0.084). Post hoc multiple comparisons of the interaction showed that in the Go condition, significant differences were identified for the comparison of low versus mid doors (P = 0.001), low versus high doors (P < 0.001) and mid versus high doors (P = 0.002). However, there were no differences in valence ratings between differently high doors in the NoGo condition. Similar results were also found for the dominance dimension of the SAM including significant main effects of the factor door height (F_2,36_ = 8.259, P = 0.001, partial η^2^ = 0.315) as well as a significant interaction of both factors (F_2,36_ = 3.974, P = 0.028, partial η^2^ = 0.181). However, there was no significant main effect of the factor imperative stimulus (F_1,18_ = 2.082, P = 0.166). The three door heights levels did not differ significantly under the NoGo condition. However, under the Go condition, significant differences were identified in low versus mid (P = 0.009), low versus high (P = 0.006) and mid versus high doors (P = 0.010).

**FIGURE 3.**
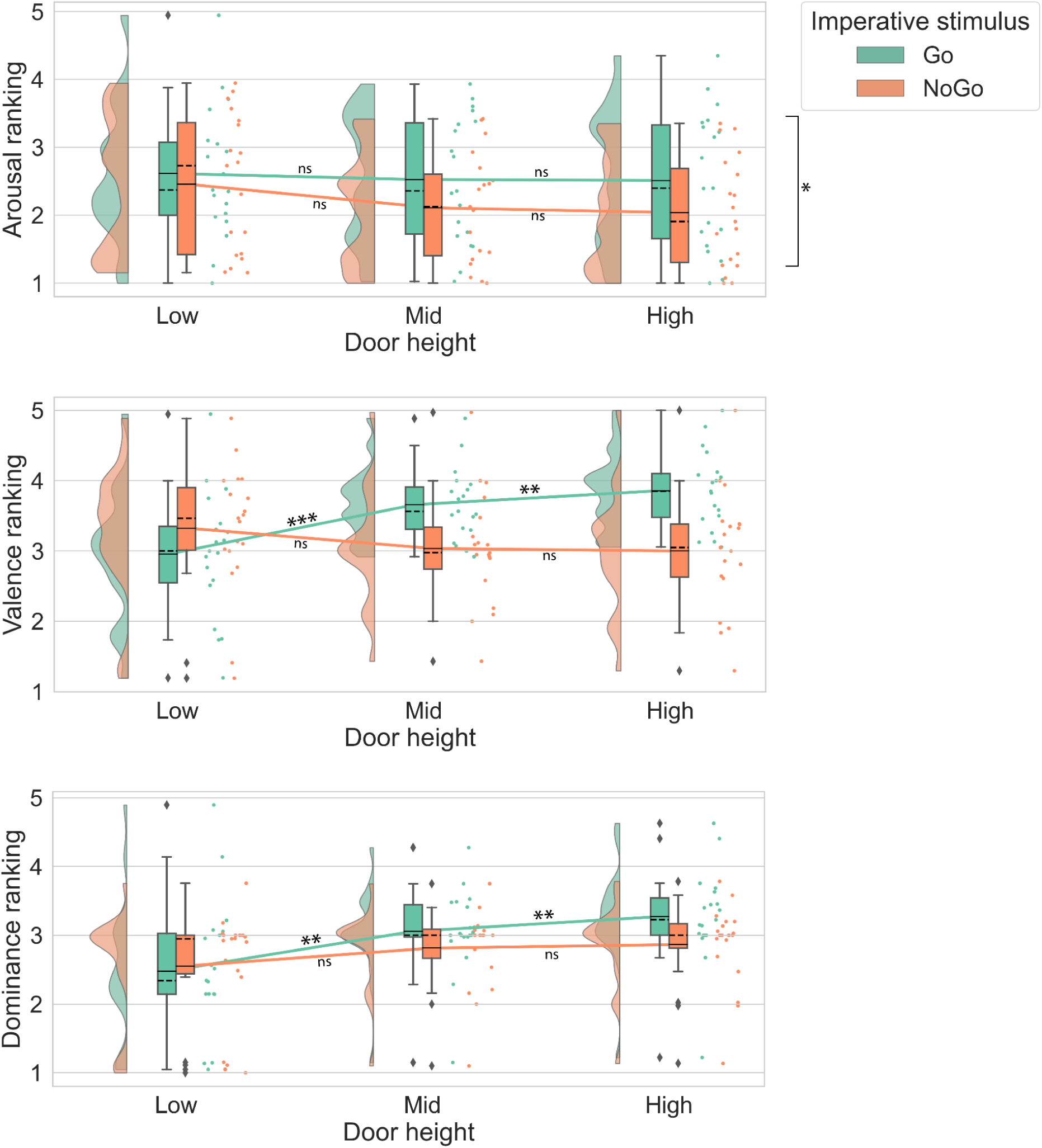
Raincloud plots displaying SAM rating for two factors: door height (low, mid and high) and imperative stimulus (Go, NoGo). The means are indicated by solid lines, and medians by dashed lines. Statistical significance was denoted by *P < 0.05, **P < 0.01 and ***P < 0.001, whereas nonsignificant results were denoted by ‘ns’.

### 3.2. Behavioral Data: Door Approaching Time

This analysis was conducted exclusively for Go trials, as these required participants to physically approach the door. The calculation was based on the time elapsed between when participants began moving out of the starting area and the moment the doors started sliding open. A one-way ANOVA with repeated measures for different door heights revealed a significant effect of door height on approach time (F_2,36_ = 3.998, P = 0.027, partial η^2^ = 0.182; figure 4). Post hoc multiple comparisons showed that significant differences were identified for the comparison of low versus high doors (P = 0.015), but no significant differences were found for the comparison of low versus mid doors (P = 0.176) and mid versus high doors (P = 0.138). It showed significantly faster approach time for the high door condition.

**FIGURE 4.**
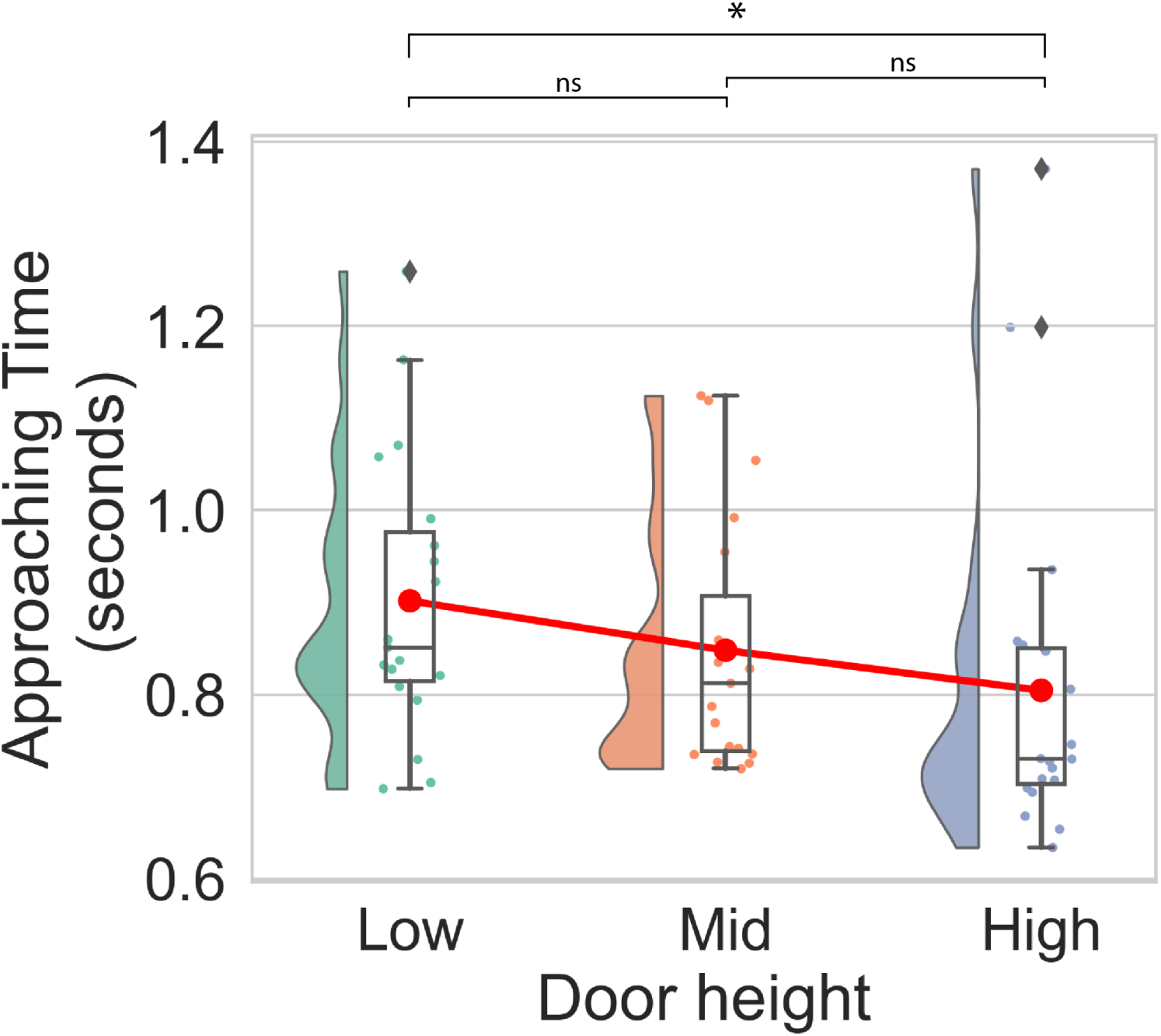
Raincloud plot of approach time for each door height condition. The means are indicated by red points and solid lines. The medians are indicated by black solid lines on each boxplot. Statistical significance was denoted as *P < 0.05, whereas nonsignificant results were denoted by ‘ns’.

### 3.3. EEG data: early visual evoked potentials

With onset of the lights that allowed participants to see the door display (i.e., lights on), the researcher identified two pronounced peaks over the occipital midline electrodes, with a first positive component around 140 ms, followed by a negative peak around 260 ms (Figure 5; all six channels shown in supplement). Similar to the findings of Djebbara et al. (2019), though not resembling a typical P1-N1 complex according to visual inspection (Figure 5), the researcher observed that at the frontal midline electrode, this pattern was inverted, with a negative component around 140 ms followed by a positive peak around 260 ms. The global 3 × 6 repeated-measures ANOVA on amplitudes with the latency of 140 ms for all 6 central midline channels revealed significant main effects for both door height (F_2,36_ = 9.383, P < 0.001, partial η^2^ = 0.343) and electrodes (F_5,90_ = 137.875, P < 0.001, partial η^2^ = 0.885), while the interaction effect was also significant (F_10,180_ = 2.123, P = 0.025, partial η^2^ = 0.106). Post hoc multiple comparisons of the interaction effect revealed significant differences in peak amplitudes at electrode Pz for the comparison of mid versus high doors (P = 0.043), at electrode POz for the comparison of low versus high doors (P = 0.012) and at electrode Oz between low and high doors (P = 0.005). However, for all anterior leads, there were no significant differences depending on affordances (Figure 6).

**FIGURE 5.**
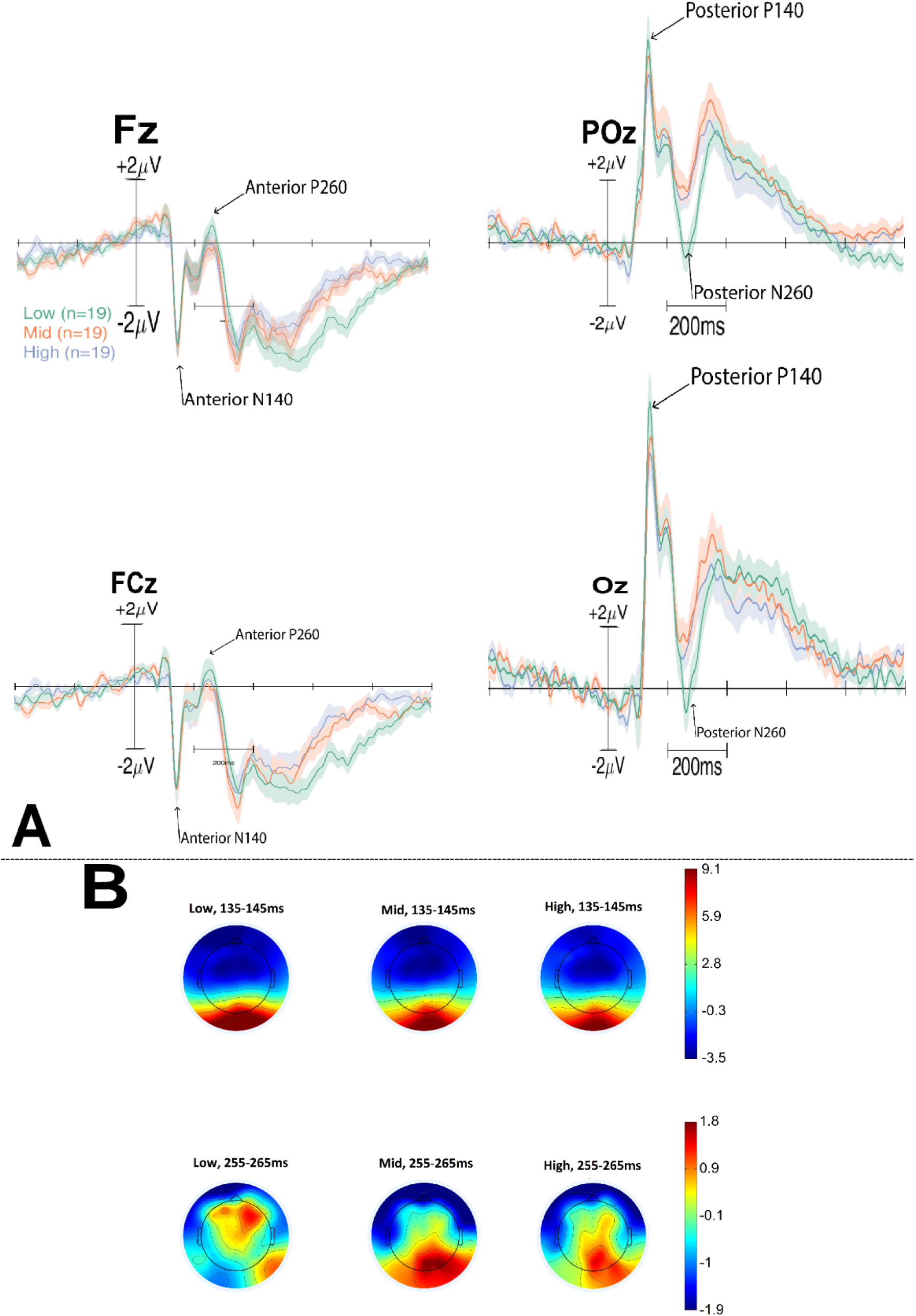
(A) Grand average event-related potentials with onset of the room display at 4 electrodes. The different door heights are colour coded with the low door condition in green, the mid door condition in red and the high door condition in purple. Prominent components (Anterior N140, P260, Posterior P140, N260) as observed over electrodes Fz, FCz, POz and Oz are indicated by arrows. (B) Scalp topographical maps depicting the differences between the door height conditions during the time window of early ERPs.

**FIGURE 6.**
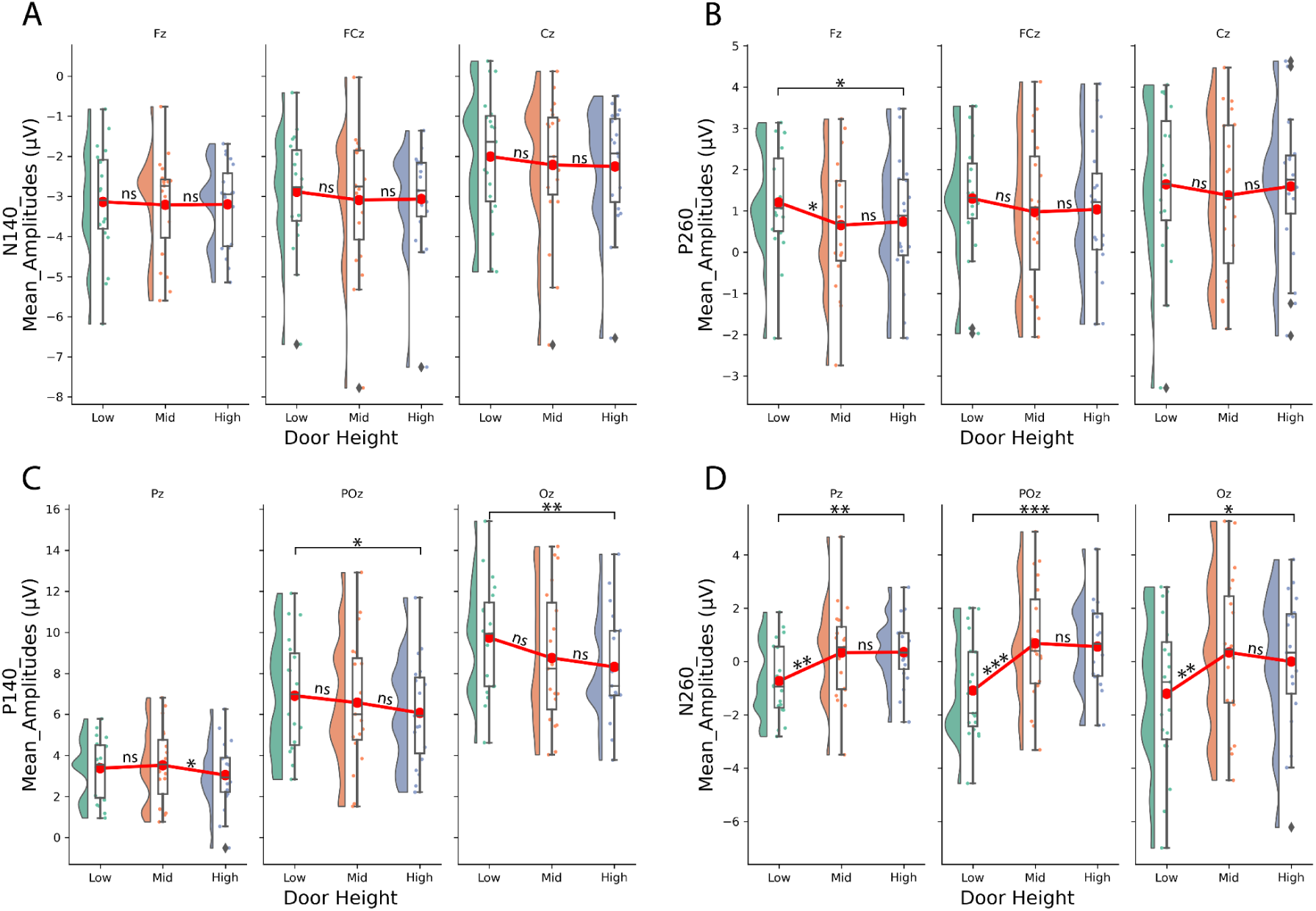
(A) Anterior N140. Raincloud plot depicting the detected mean amplitude of the negative peak in the time-locked ’lights on’ event, within the 90–190 ms time range, for electrodes Fz, FCz, and Cz. (B) Anterior P260. Raincloud plot depicting the detected mean amplitude of the positive peak in the time-locked ’lights on’ event, within the 210–310 ms time range, for electrodes Fz, FCz, and Cz. (C) Posterior P140. Raincloud plot depicting the detected mean amplitude of the positive peak in the time-locked ’lights on’ event, within the 90–190 ms time range, for electrodes Pz, POz, and Oz. (D) Posterior N260. Raincloud plot depicting the detected mean amplitude of the negative peak in the time-locked ’lights on’ event, within the 210–310 ms time range, for electrodes Pz, POz, and Oz. The means are indicated by red points and solid lines. The medians are indicated by black solid lines on each boxplot. The low condition is in green, the mid condition is in red, and the high condition is in purple. Statistical significance was denoted by *P < 0.05, **P < 0.01 and ***P < 0.001, whereas non-significant results were denoted by ‘ns’.

The global 3 × 6 repeated-measures ANOVA on amplitudes with the latency of 260 ms for all 6 central midline channels revealed significant main effects for both door height (F_2,36_ = 10.678, P < 0.001, partial η^2^ = 0.372) and electrodes (F_5,90_ = 3.406, P = 0.007, partial η^2^ = 0.159). And the interaction effect was also significant (F_10,180_ = 6.772, P < 0.001, partial η^2^ = 0.273). Post hoc multiple comparisons of the interaction effect revealed significant differences in peak amplitudes at electrode Pz for the comparison of low versus mid doors (P = 0.006) and of low versus high doors (P = 0.003), at electrode POz for the comparison of low versus mid doors (P < 0.001) and of low versus high doors (P < 0.001) and at electrode Oz between low and mid doors (P = 0.005) and between low and high doors (P = 0.011). However, for all anterior leads, there was only an affordance effect for the electrode Fz. Significant differences were identified at this channel for the comparison of low versus mid doors (P = 0.049) and low versus high doors (P = 0.020), but no significant differences were found for the comparison of mid versus high doors (P = 0.712).

### 3.4. Motor-related ERPs: PINV

The PINV component was observed at 1100–1400 ms after the imperative stimulus (Go/NoGo) and varied as a function of the environmental affordances (figure 7). A global 2 × 3 × 6 factorial repeated-measures ANOVA was computed to analyze the PINV using the factors imperative stimulus (Go and NoGo), door height (low, mid, and high), and electrode location (Fz, FCz, Cz, Pz, POz, and Oz). The ANOVA revealed a significant main effect for the factor imperative stimulus (F_1,18_= 5.559, P = 0.030, partial η^2^ = 0.236) and a significant interaction effect of imperative stimulus × door height × electrodes (F_10,180_ = 2.045, P = 0.031, partial η^2^ = 0.102). The researcher did not find a significant main effect for the factor door height (F_2,36_ = 2.722, P = 0.079).

**FIGURE 7.**
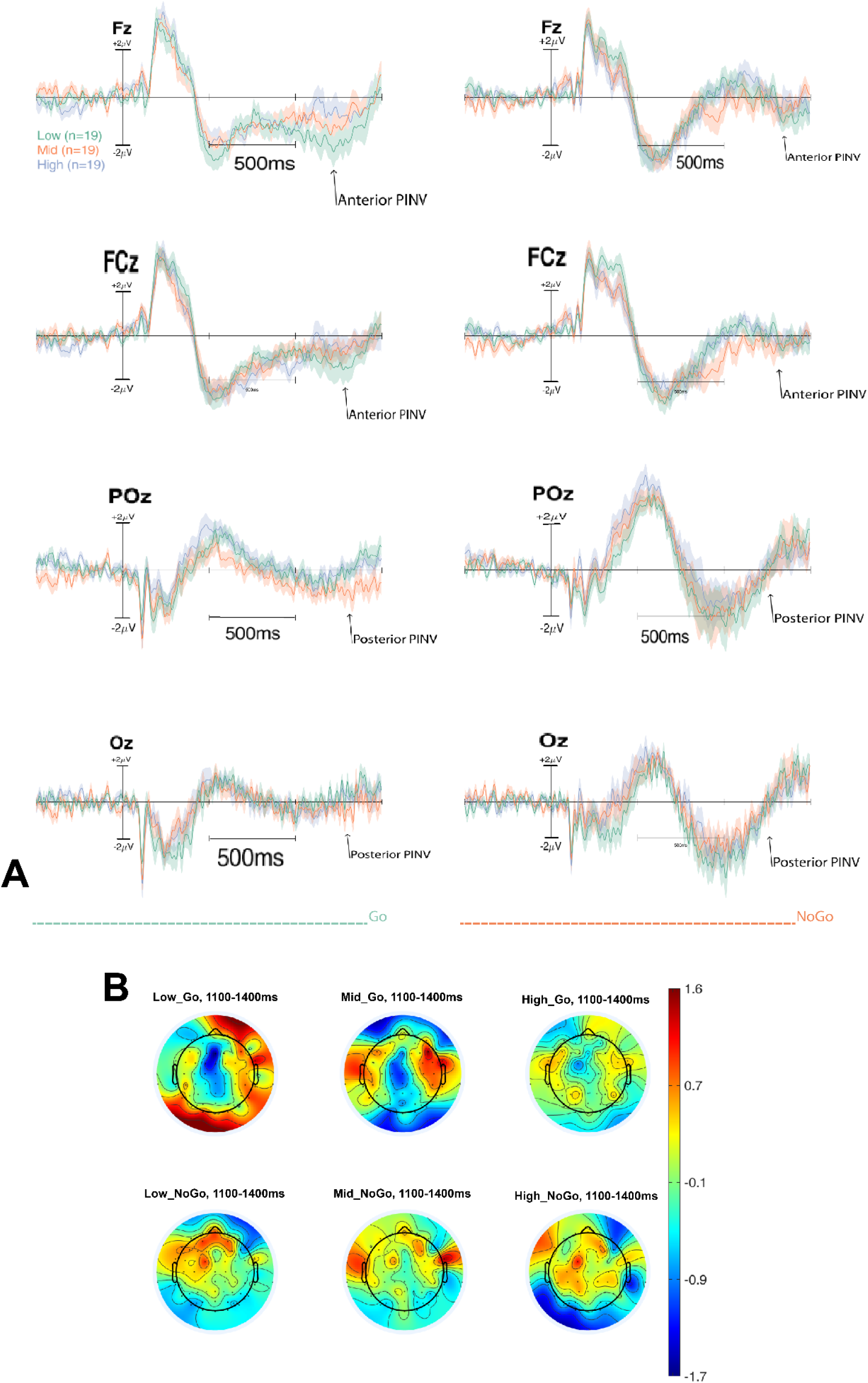
(A) Eight time-locked event-related potentials (ERPs) from electrodes Fz, FCz, POz and Oz at the onset of imperative stimulus (Go, NoGo). The low condition is in green, the mid condition is in red, and the high condition is in purple. The plots on the left column showed the ERPs under the Go condition, whereas the plots on the right column showed the ERPs under the NoGo condition. PINV components were marked with arrows. (B) Scalp topographical maps depicting the differences between the door height conditions and the differences between Go and NoGo trials during the time window of PINV.

Interaction effects showed significant differences of transitional affordances emerged only under the Go condition (figure 8). Post hoc comparisons of this three-way interaction revealed significant differences at electrode Fz for the comparison of low versus mid doors (P = 0.019) and of low versus high doors (P = 0.038). No significant differences were found at electrode POz between low and mid doors (P = 0.089) or between mid and high doors (P = 0.062) nor at electrode Oz for the comparison of low versus mid doors (P = 0.080).

**FIGURE 8.**
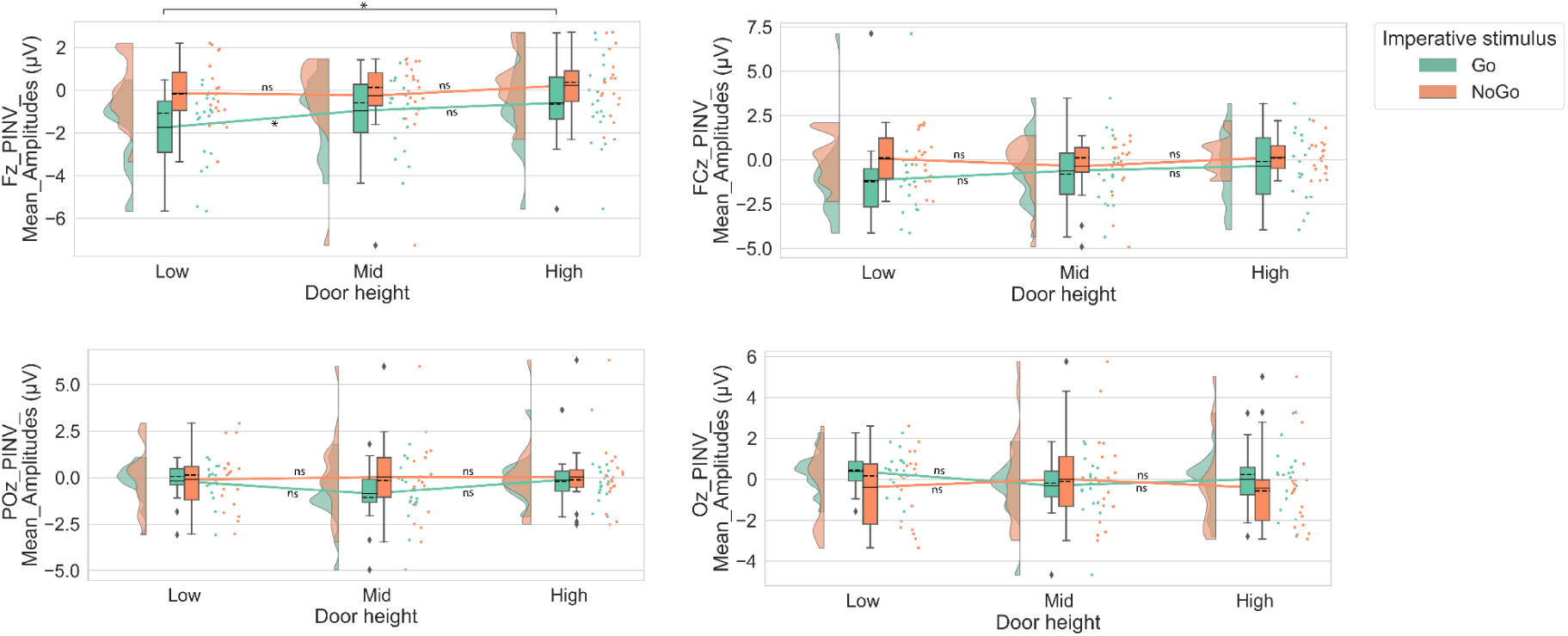
Raincloud plots of mean amplitude of negative development in the time-locked event of Go/NoGo in the time range of 1100–1400 ms for Fz, FCz, POz, and Oz. The means are indicated by solid lines, and medians by dashed lines. Statistical significance was denoted by *P < 0.05, whereas nonsignificant results were denoted by ‘ns’.

## 4. Discussion

In view of the conceptual tension between micro and macro affordances (also known as environmental affordances), the main goal of this study was to investigate whether an environmental affordance can be manipulated by an intrinsic setting and whether varying levels of affordance manipulation could prime agents’ experience, motor interaction and perception. By providing this paradigm extension (macro-intrinsic affordances), not only does the present study wish to mitigate the aforementioned tension but also to provide empirical support for the need to link ecological psychology and neuro-architecture in experimental protocols (for a theoretical review see S. Wang et al., 2022). If this were the case, then perceptual processes would covary with the environmental-intrinsic affordances, leading to subjective rating and behavioral changes, and that both early ERPs and the later MRCPs would vary as functions of affordances. Focusing on this research target, in order to operationally define environmental affordance in an intrinsic manner, the present paradigm adapted the task of Djebbara et al. (2019) by defining transitional affordance as the body-door height ratios. To investigate the research question, experimenters collected subjective, behavioral, and electrophysiological data, with a focus on electrophysiology.

### 4.1. SAM Ratings

The analysis of the SAM ratings for valence and dominance consistently revealed significant differences dependent on door height among Go trials but no differences across NoGo trials for any rating. However, for the arousal dimension, only two main effects were observed, with participants reporting greater arousal for low doors compared to mid and high doors, as well as greater arousal for Go trials than NoGo trials. Post-interviews with some participants revealed that they experienced greater emotional fluctuations after viewing the low door, as squeezing their bodies through it required considerable effort. Regarding the interaction effect of Go/NoGo × height, varying door sizes for Go trials yielded differences for dominance, with low doors less dominating than Mid doors and even less dominating than high doors. The present research discovered a similar pattern for the dimension of valence. Post-interviews with some participants revealed that they were happier when they viewed a green mid or green high door than a green low door, because they can easily pass through the transition and get their bonus. Taken together, when the participants knew that they would interact with doors of different transition affordances (Go trials), data can present a distinct impact of action affective ratings of an environment. Nevertheless, the affordance effect should be interpreted cautiously, as the subjective data may have been influenced by various factors beyond affordances, such as differing motivation levels from distinct reward settings (monetary compensation or accredited course hours), differences in trial durations, physical exertion, and individual variations in introspective ability.

### 4.2. Approaching Time

In contrast to the potential confounders associated with subjective data, performance data may provide a more reliable basis for interpreting the impact of affordances on behavior, given the relative consistency of motor trajectories and approach strategies. In the present study, the low door condition allowed participants to pass through, albeit at a significant physical cost, and the mid door also afforded opportunities for body folding, causing participants to maintain a relatively high level of enthusiasm and concentration throughout the trials. The time it took participants to reach the door after the onset of the imperative stimulus varied according to the environmental affordance. Although the current environmental affordance was scaled intrinsically, the approach time differed significantly across conditions.

Participants approached the high doors significantly faster than the mid and low doors, respectively, while there was no significant difference between the latter two. The high door allowed easy passage without imposing notable demands on motor planning or execution. In contrast, the mid-height door, set at 95% of individual body heights, prompted motor anticipation, as participants evaluated the passability and slightly adjusted their body posture while approaching. Although the low door was set at 60% of body height, participants could still pass, adopting a squat or duck-walking strategy to do so. In this sense, both the mid and low doors may have delayed approaching time due to increased processing demands, though to different extents. Although the difference in physical demands between the low and mid doors did not reach statistical significance in the behavioral data, it is promising that cortical potentials related to environmental affordances could provide more robust data. EEG data, being less influenced by subjective interpretation and external factors, may offer a more objective way to interpret the sensorimotor dynamics and proprioceptive mechanisms underlying the observation of approaching time and motor trajectory.

### 4.3. Early evoked potentials

To examine whether the environmental-intrinsic affordance can be reflected by brain dynamics and whether underlying sensorimotor brain coordination differs according to different sensorimotor stages, experimenters recorded both early visual evoked potentials and motor-related potentials. Building on the fast coupling between environmental features and sensorimotor responses (referred to as the “EF-SMR relation”; see Djebbara et al., 2022) and neural reuse theory (Anderson, 2010), the functional coupling between environmental affordances and associated sensorimotor responses should be activated automatically within the early sensorimotor stage, potentially involving a coordination pattern between different brain regions. Guided by this framework, the present research focused on early evoked potentials across six central midline channels and conducted a global ANOVA and topographical mapping, and expected to find differences in the stimulus-locked ERPs under different affordance conditions. The current study was cautious with interpreting main effects since polarity reversals may potentially cancel each other out and thus focused on the interaction effects and post hoc multiple comparisons. As illustrated in the result section, the present research found significant differences in amplitudes of the visually evoked P140 and N260 component over the posterior channels varying with the affordance of the transition. In addition, the researcher also found a difference in amplitude of the visually evoked P260 component at the channel Fz after onset of the door display. Notably, the global ANOVA results and topographic patterns differed between posterior and anterior channels, with the posterior area more pronounced activation. In short, these findings indicated that transitional affordances can be manipulated by an intrinsic setting and then varying levels of affordance manipulation could prime cortical potentials in the early sensorimotor time window, following an anteroposterior gradient.

The current early evoked ERPs results, in terms of the level of peak latencies, replicate the previous finding of our team (Djebbara et al., 2019; S. Wang et al., 2024). Consistent with the hypothesis, the results demonstrate an association between environmental affordances and early visual-evoked potentials. The researcher did not observe a typical P1-N1 complex, either through visual inspection (Figure 5) or statistical analysis (Figure 6). Based on the peak latency information and prior research, this paper referred to the posterior P140 and N260 as posterior P1 and N2 (Patel & Azzam, 2005), respectively, and the anterior P260 as anterior P2 (Hakkarainen et al., 2012; Yu et al., 2021). Regarding the visual evoked potentials at occipital sites, prior research has shown that the P1 represents a facilitation of early sensory processing for stimuli occurring in the attended location (Luck et al., 1990). The N2 component represents attentional selection of relevant information, voluntary attention and the capability of inhibiting distraction (Luck & Hillyard, 1994; Patel & Azzam, 2005; Schneider et al., 2012; Y. Wang et al., 2010), as well as motor anticipation or prospective movement (Kourtis et al., 2012). As we interpreted in the recent article (S. Wang et al., 2024), early perception processes of environmental affordances were associated with increased attentional processing. After the same stimulus onset, but with extended latencies, more voluntary attention was involved. This involvement might be attributed to an increased coordination of other psychological processes. While the posterior N2 component and the anterior P2 component have similar latencies, the anterior P2 reflects task-relevant motor planning (Yu et al., 2021). This suggests that even during the early sensorimotor stage, coordination between sensory and motor activities occurs, aligning with the EF-SMR relation (Djebbara et al., 2022) and neural reuse theory (Anderson, 2010).

Taken together, in line with the findings of Djebbara et al. (2019), the current work found that early perceptual processes covary with environmental affordances. This covariation mechanism was stable irrespective of whether the affordance was scaled intrinsically or not. In addition, this covariation was found at both posterior and anterior regions, with more pronounced activation in the posterior area. The former finding reflected that the rapid perception processes related to environmental affordance were shared across different brain regions, while the latter indicated that this sharing was not entirely uniform, but followed an anteroposterior gradient. In other words, the division of brain processing underlying the environmental affordances could be differentiated but shall not be separated. This finding was in line with the account of neural reuse (Anderson, 2010).

### 4.4. Motor-related potentials: PINV

After the onset of imperative stimulus, a prominent PINV component was observed with a latency between 1100 to 1400 ms. The global three-way interaction effect among the factors imperative stimulus, door height, and electrodes was observed, indicating a potential brain localization of PINV. The present study found differences in frontal, parieto-occipital and occipital areas, with the PINV amplitudes being most pronounced at Fz. At this electrode, a two-way interaction effect between the factors imperative stimulus and door height was observed. Only in cases of Go did the researcher observe a pronounced difference in the PINV component, which varied with the environmental affordances. And there were no significant differences in the PINV component in cases of NoGo. Thus in the current protocol, the environmental affordance can be manipulated by an intrinsic setting and then varying levels of affordance manipulation could prime motor-related brain dynamics. The negative affordance emerged when architectural transitions constrained a walking agent’s body range to a certain extent. In this case, the extent was predefined as a constant geometric ratio (60%) between door and body height, regardless of the absolute sizes of the agents’ heights or the door heights.

Regarding the psychological meaning of PINV, studies have shown that it is a suitable indicator of information processing during lack of control over aversive events and generally PINV was more pronounced at the frontal (Fz) than at the precentral (Cz) and parietal (Pz) recording sites (Diener, Struve, et al., 2009; Elbert et al., 1982). In the present case, the “aversive events” can be regarded as negative affordances (low doors) as participants had to adopt a squat or duck-walking strategy so as to pass through the small transition. This limitation likely caused their lower valence scores and delayed approaching time. And the lower dominating ranking of the low door condition reflected a sense of loss of control when they had to pay a lot of physical effort.

Regarding the above post hoc comparisons of the two-way interaction between the factors imperative stimulus and door height, it reflected that PINV was evoked by the imminent motor execution and also reshaped by the anticipated motor trajectory (in line with Djebbara et al., 2019). According to our environmental experience, our action was executed and extended based on given perceptions, forming corresponding motor trajectories. Interestingly, the PINV data revealed a notable distinction: first, neural activity depended on whether the door was perceived as easily passable or not. Second, this dependency was found only when the person knew that he or she would need to interact with the door. It thus supported the view that the world of perception is dynamically reshaped by the possibilities of imminent action, namely environmental affordances. However, homogeneous PINV amplitudes in the “NoGo” condition should not be interpreted as action preparation. When the door color turned to “red”, participants did not need to interact with the transition but instead simply turned back to answer the questionnaire. Therefore, in this case, the brain dynamics data might not be attributed to an anticipated motor trajectory or proprioceptive information of varying transition affordances. Rather, it likely corresponded to a uniform and straightforward body-turning movement.

Regarding the modulation of the factor electrodes in the three-way interaction effect, similar to a previous study (Elbert et al., 1982), the researcher observed differences in frontal, parietal-occipital and occipital areas, with the PINV waveform most pronounced at Fz. Combining the global ANOVA results and topographic patterns, the current work observed a reversal of the anteroposterior gradient compared to the early ERPs, which suggests a potential modulation originating from sensorimotor stages. It was in line with the neural reuse theory and the account of the sensorimotor schemes (Anderson, 2010; Djebbara et al., 2022; Paolo et al., 2017). While this opinion was based on a qualitative comparison, conducting an EEG connectivity analysis in the next step would be valuable to further explore time-varying brain coordination, as it echoes our recent findings on time-varying automaticity of affordance perception (S. Wang et al., 2024).

## 5. Conclusion

Using mobile brain/body imaging and an intrinsic paradigm, the present study provides a synthesis-like perspective that harmonizes the conceptual tension between micro-affordances and macro/environmental affordances. Focusing on virtual transitions with differing body-door height ratios, our results indicate that environmental affordances can be manipulated through an ecological principle called intrinsic measurement. The perception of environmental affordances was thus based on intrinsic optical information for the relationship between environmental properties and capacities of the observer’s own action system. When architectural transitions constrain an agent’s body range to a certain extent, negative affordances emerge, impacting experience, behavior, and perception processes. This perception is body-scaled, action-oriented, and thus enactive and dynamic. Initial perception provides the foundation for subsequent spatial exploration and motor anticipation, while our environmental perception is dynamically reshaped by the possibilities of imminent action. First, the current study extended the intrinsic paradigm from the experimental protocol of micro-affordances to macro-affordances, demonstrating that human-environment interactions are inherently linked to the agent’s body scale. Second, this paradigm extension thus offers a promising tie between ecological psychology and neuro-architecture in experimental protocols, as advocated for in our prior theoretical review (S. Wang et al., 2022).

## Statement

During the preparation of this work the author used the AI-based language model ChatGPT (version 3.5) for text refinement and enhancement during the writing process. After using this tool, the author reviewed and edited the content as needed and took full responsibility for the content of the publication.

## Data and codes availability

Codes (EEG preprocessing, ERPs epoching, ERPs calculation and data visualization) were uploaded to github (https://github.com/Feng-xiaoming/Architectual-affordances-door-height). The data files are publicly available through an OSF repository (DOI: 10.17605/OSF.IO/42DZS).

## CRediT authorship contribution statement

**Sheng Wang:** Writing – review & editing, Writing – original draft, Visualization, Methodology, Investigation, Formal analysis, Data curation, Conceptualization.

## Acknowledgements

The author would like to thank his supervisor Klaus Gramann for his valuable comments and advice on article writing, team engineer Benjamin Paulisch for his Unity programming, colleague Christopher Hilton for his advice regarding statistical analysis, student assistant Luca Strasser for his lab assistance, his academic peer Zhenxing Hong and student assistant Magda Biadala for their assistance in enhancing the graphic aesthetics of the present study.

## Reference

1. Altomare, E. C., Committeri, G., Di Matteo, R., Capotosto, P., & Tosoni, A. (2021). Automatic coding of environmental distance for walking-related locomotion in the foot-related sensory-motor system: A TMS study on macro-affordances. Neuropsychologia, 150, 107696. 10.1016/j.neuropsychologia.2020.107696

2. Anderson, M. L. (2010). Neural reuse: A fundamental organizational principle of the brain. Behavioral and Brain Sciences, 33(4), 245–266. 10.1017/S0140525X10000853

3. Aron, A. R., & Poldrack, R. A. (2005). The Cognitive Neuroscience of Response Inhibition: Relevance for Genetic Research in Attention-Deficit/Hyperactivity Disorder. Biological Psychiatry, 57(11), 1285–1292. 10.1016/j.biopsych.2004.10.026

4. Bonner, M. F., & Epstein, R. A. (2017). Coding of navigational affordances in the human visual system. Proceedings of the National Academy of Sciences, 114(18), 4793–4798. 10.1073/pnas.1618228114

5. Bozzacchi, C., Giusti, M. A., Pitzalis, S., Spinelli, D., & Di Russo, F. (2012). Awareness affects motor planning for goal-oriented actions. Biological Psychology, 89(2), 503–514. 10.1016/j.biopsycho.2011.12.020

6. Bozzacchi, C., Spinelli, D., Pitzalis, S., Giusti, M. A., & Di Russo, F. (2015). I know what I will see: Action-specific motor preparation activity in a passive observation task. Social Cognitive and Affective Neuroscience, 10(6), 783–789. 10.1093/scan/nsu115

7. Bradley, M. M., & Lang, P. J. (1994). Measuring emotion: The self-assessment manikin and the semantic differential. Journal of Behavior Therapy and Experimental Psychiatry, 25(1), 49–59. 10.1016/0005-7916(94)90063-9

8. Casement, M. D., Shestyuk, A. Y., Best, J. L., Casas, B. R., Glezer, A., Segundo, M. A., & Deldin, P. J. (2008). Anticipation of affect in dysthymia: Behavioral and neurophysiological indicators. Biological Psychology, 77(2), 197–204. 10.1016/j.biopsycho.2007.10.007

9. Chemero, A. (2003). An Outline of a Theory of Affordances. In How Shall Affordances Be Refined? Routledge.

10. Chemero, A. (2013). Radical Embodied Cognitive Science. Review of General Psychology, 17(2), 145–150. 10.1037/a0032923

11. Costantini, M., Ambrosini, E., Tieri, G., Sinigaglia, C., & Committeri, G. (2010). Where does an object trigger an action? An investigation about affordances in space. Experimental Brain Research, 207(1), 95–103. 10.1007/s00221-010-2435-8

12. Delorme, A., & Makeig, S. (2004). EEGLAB: An open source toolbox for analysis of single-trial EEG dynamics including independent component analysis. Journal of Neuroscience Methods, 134(1), 9–21. 10.1016/j.jneumeth.2003.10.009

13. Derbyshire, N., Ellis, R., & Tucker, M. (2006). The potentiation of two components of the reach-to-grasp action during object categorisation in visual memory. Acta Psychologica, 122(1), 74–98. 10.1016/j.actpsy.2005.10.004

14. Di Marco, S., Tosoni, A., Altomare, E. C., Ferretti, G., Perrucci, M. G., & Committeri, G. (2019). Walking-related locomotion is facilitated by the perception of distant targets in the extrapersonal space. Scientific Reports, 9(1), 9884. 10.1038/s41598-019-46384-5

15. Diener, C., Kuehner, C., Brusniak, W., Struve, M., & Flor, H. (2009). Effects of stressor controllability on psychophysiological, cognitive and behavioural responses in patients with major depression and dysthymia. Psychological Medicine, 39(1), 77–86. 10.1017/S0033291708003437

16. Diener, C., Struve, M., Balz, N., Kuehner, C., & Flor, H. (2009). Exposure to uncontrollable stress and the postimperative negative variation (PINV): Prior control matters. Biological Psychology, 80(2), 189–195. 10.1016/j.biopsycho.2008.09.002

17. Djebbara, Z., Fich, L. B., Petrini, L., & Gramann, K. (2019). Sensorimotor brain dynamics reflect architectural affordances. Proceedings of the National Academy of Sciences, 116(29), 14769–14778. 10.1073/pnas.1900648116

18. Djebbara, Z., Jensen, O. B., Parada, F. J., & Gramann, K. (2022). Neuroscience and architecture: Modulating behavior through sensorimotor responses to the built environment. Neuroscience & Biobehavioral Reviews, 138, 104715. 10.1016/j.neubiorev.2022.104715

19. Elbert, T., Rockstroh, B., Lutzenberger, W., & Birbaumer, N. (1982a). Slow brain potentials after withdrawal of control. Archiv Für Psychiatrie Und Nervenkrankheiten, 232(3), 201–214. 10.1007/BF02141781

20. Elbert, T., Rockstroh, B., Lutzenberger, W., & Birbaumer, N. (1982b). Slow brain potentials after withdrawal of control. Archiv Für Psychiatrie Und Nervenkrankheiten, 232(3), 201–214. 10.1007/BF02141781

21. Ellis, R., & Tucker, M. (2000). Micro-affordance: The potentiation of components of action by seen objects. British Journal of Psychology, 91(4), 451–471. 10.1348/000712600161934

22. Feng, X., Xu, S., Li, Y., & Liu, J. (2024). Body size as a metric for the affordable world. eLife, 12, RP90583. 10.7554/eLife.90583

23. Ferretti, G. (2016). Pictures, action properties and motor related effects. Synthese, 193(12), 3787–3817. 10.1007/s11229-016-1097-x

24. Ferretti, G. (2021). A distinction concerning vision-for-action and affordance perception. Consciousness and Cognition, 87, 103028. 10.1016/j.concog.2020.103028

25. Gibson, J. J. (1966). The senses considered as perceptual systems. Houghton Mifflin. Gibson, J. J. (2014). The Ecological Approach to Visual Perception: Classic Edition. Psychology Press. 10.4324/9781315740218

26. Goslin, J., Dixon, T., Fischer, M. H., Cangelosi, A., & Ellis, R. (2012). Electrophysiological Examination of Embodiment in Vision and Action. Psychological Science, 23(2), 152–157. 10.1177/0956797611429578

27. Gregorians, L., & Spiers, H. J. (2022). Affordances for Spatial Navigation. In Z. Djebbara (Ed.), Affordances in Everyday Life: A Multidisciplinary Collection of Essays (pp. 99–112). Springer International Publishing. 10.1007/978-3-031-08629-8_10

28. Hakkarainen, E., Pirilä, S., Kaartinen, J., & Meere, J. J. van der. (2012). Stimulus Evaluation, Event Preparation, and Motor Action Planning in Young Patients With Mild Spastic Cerebral Palsy: An Event-Related Brain Potential Study. Journal of Child Neurology, 27(4), 465–470. 10.1177/0883073811420150

29. Harel, A., Nador, J. D., Bonner, M. F., & Epstein, R. A. (2022). Early Electrophysiological Markers of Navigational Affordances in Scenes. Journal of Cognitive Neuroscience, 34(3), 397–410. 10.1162/jocn_a_01810

30. Jungnickel, E., Gehrke, L., Klug, M., & Gramann, K. (2019). Chapter 10—MoBI—Mobile Brain/Body Imaging. In H. Ayaz & F. Dehais (Eds.), Neuroergonomics (pp. 59–63). Academic Press. 10.1016/B978-0-12-811926-6.00010-5

31. Klug, M., & Gramann, K. (2021). Identifying key factors for improving ICA-based decomposition of EEG data in mobile and stationary experiments. European Journal of Neuroscience, 54(12), 8406–8420. 10.1111/ejn.14992

32. Klug, M., Jeung, S., Wunderlich, A., Gehrke, L., Protzak, J., Djebbara, Z., Argubi-Wollesen, A., Wollesen, B., & Gramann, K. (2022). The BeMoBIL Pipeline for automated analyses of multimodal mobile brain and body imaging data (p. 2022.09.29.510051). bioRxiv. 10.1101/2022.09.29.510051

33. Klug, M., & Kloosterman, N. A. (2022). Zapline-plus: A Zapline extension for automatic and adaptive removal of frequency-specific noise artifacts in M/EEG. Human Brain Mapping, 43(9), 2743–2758. 10.1002/hbm.25832

34. Kothe, C., Medine, D., Boulay, C., Grivich, M., & Stenner, T. (2014). *Lab streaming layer*.

35. Kourtis, D., Sebanz, N., & Knoblich, G. (2012). EEG correlates of Fitts’s law during preparation for action. Psychological Research, 76(4), 514–524. 10.1007/s00426-012-0418-z

36. Luck, S. J., Heinze, H. J., Mangun, G. R., & Hillyard, S. A. (1990). Visual event-related potentials index focused attention within bilateral stimulus arrays. II. Functional dissociation of P1 and N1 components. Electroencephalography and Clinical Neurophysiology, 75(6), 528–542. 10.1016/0013-4694(90)90139-B

37. Luck, S. J., & Hillyard, S. A. (1994). Electrophysiological correlates of feature analysis during visual search. Psychophysiology, 31(3), 291–308. 10.1111/j.1469-8986.1994.tb02218.x

38. Makeig, S., Gramann, K., Jung, T.-P., Sejnowski, T. J., & Poizner, H. (2009). Linking brain, mind and behavior. International Journal of Psychophysiology, 73(2), 95–100. 10.1016/j.ijpsycho.2008.11.008

39. Oostenveld, R., & Praamstra, P. (2001). The five percent electrode system for high-resolution EEG and ERP measurements. Clinical Neurophysiology, 112(4), 713–719. 10.1016/S1388-2457(00)00527-7

40. Palmer, J. A., Kreutz-Delgado, K., & Makeig, S. (2011). *AMICA: An Adaptive Mixture of Independent Component Analyzers with Shared Components*.

41. Paolo, E. D., Buhrmann, T., & Barandiaran, X. (2017). Sensorimotor Life: An enactive proposal. Oxford University Press.

42. Patel, S. H., & Azzam, P. N. (2005). Characterization of N200 and P300: Selected Studies of the Event-Related Potential. International Journal of Medical Sciences, 2(4), 147–154.

43. Pezzulo, G., & Cisek, P. (2016). Navigating the Affordance Landscape: Feedback Control as a Process Model of Behavior and Cognition. Trends in Cognitive Sciences, 20(6), 414–424. 10.1016/j.tics.2016.03.013

44. Pion-Tonachini, L., Kreutz-Delgado, K., & Makeig, S. (2019). ICLabel: An automated electroencephalographic independent component classifier, dataset, and website. NeuroImage, 198, 181–197. 10.1016/j.neuroimage.2019.05.026

45. Rowe, P. (2018). *The temporal nature of affordance: An investigation using EEG and TMS* [Doctoral, City, University of London]. https://openaccess.city.ac.uk/id/eprint/20554/

46. Schneider, D., Beste, C., & Wascher, E. (2012). On the time course of bottom-up and top-down processes in selective visual attention: An EEG study. Psychophysiology, 49(11), 1660–1671. 10.1111/j.1469-8986.2012.01462.x

47. Tosoni, A., Altomare, E. C., Brunetti, M., Croce, P., Zappasodi, F., & Committeri, G. (2021). Sensory-Motor Modulations of EEG Event-Related Potentials Reflect Walking-Related Macro-Affordances. Brain Sciences, 11(11), Article 11. 10.3390/brainsci11111506

48. Tosoni, A., Altomare, E. C., Perrucci, M. G., Committeri, G., & Di Matteo, R. (2023). Foot-related/walking macro-affordances are implicitly activated and preferentially guided by the framing distance of the environmental layout. Psychological Research, 87(3), 787–799. 10.1007/s00426-022-01692-w

49. Tucker, M., & Ellis, R. (1998). On the relations between seen objects and components of potential actions. Journal of Experimental Psychology: Human Perception and Performance, 24(3), 830–846. 10.1037/0096-1523.24.3.830

50. Tucker, M., & Ellis, R. (2001). The potentiation of grasp types during visual object categorization. Visual Cognition, 8(6), 769–800. 10.1080/13506280042000144

51. Tucker, M., & Ellis, R. (2004). Action priming by briefly presented objects. Acta Psychologica, 116(2), 185–203. 10.1016/j.actpsy.2004.01.004

52. Verleger, R., Wascher, E., Arolt, V., Daase, C., Strohm, A., & Kömpf, D. (1999). Slow EEG potentials (contingent negative variation and post-imperative negative variation) in schizophrenia: Their association to the present state and to Parkinsonian medication effects. Clinical Neurophysiology, 110(7), 1175–1192. 10.1016/S1388-2457(99)00023-1

53. Wang, S., Djebbara, Z., Sanches de Oliveira, G., & Gramann, K. (2024). Human brain dynamics dissociate early perceptual and late motor-related stages of affordance processing. European Journal of Neuroscience, 60(4), 4639–4660. 10.1111/ejn.16461

54. Wang, S., Sanches de Oliveira, G., Djebbara, Z., & Gramann, K. (2022). The Embodiment of Architectural Experience: A Methodological Perspective on Neuro-Architecture. Frontiers in Human Neuroscience, 16. 10.3389/fnhum.2022.833528

55. Wang, Y., Wu, J., Fu, S., & Luo, Y. (2010). Orienting and Focusing in Voluntary and Involuntary Visuospatial Attention Conditions. Journal of Psychophysiology, 24(3), 198–209. 10.1027/0269-8803/a000010

56. Warren, W. H. (1984). Perceiving affordances: Visual guidance of stair climbing. Journal of Experimental Psychology: Human Perception and Performance, 10(5), 683–703. 10.1037/0096-1523.10.5.683

57. Yu, L., Schack, T., & Koester, D. (2021). Coordinating Initial and Final Action Goals in Planning Grasp-to-Rotate Movements: An ERP Study. Neuroscience, 459, 70–84. 10.1016/j.neuroscience.2021.01.033

58. Zipoli Caiani, S. (2014). Extending the notion of affordance. Phenomenology and the Cognitive Sciences, 13(2), 275–293. 10.1007/s11097-013-9295-1

