## Supplement for "Manipulating Environmental Affordances via Intrinsic Measurement: Evidence from Sensorimotor Brain Dynamics"

This PDF file includes:

Fig. S1: ERP plots for early ERPs at 6 electrodes (Fz, FCz, Cz, Pz, POz, Oz)

Fig. S2: ERP plots for PINV at 6 electrodes (Fz, FCz, Cz, Pz, POz, Oz)

Fig. S3: Low door green

Fig. S4: Low door grey

Fig. S5: Low door red

Fig. S6: Mid door green

Fig. S7: Mid door grey

Fig. S8: Mid door red

Fig. S9: High door green

Fig. S10: High door grey

Fig. S11: High door red

Fig. S12: Second room

Fig. S13: Starting room

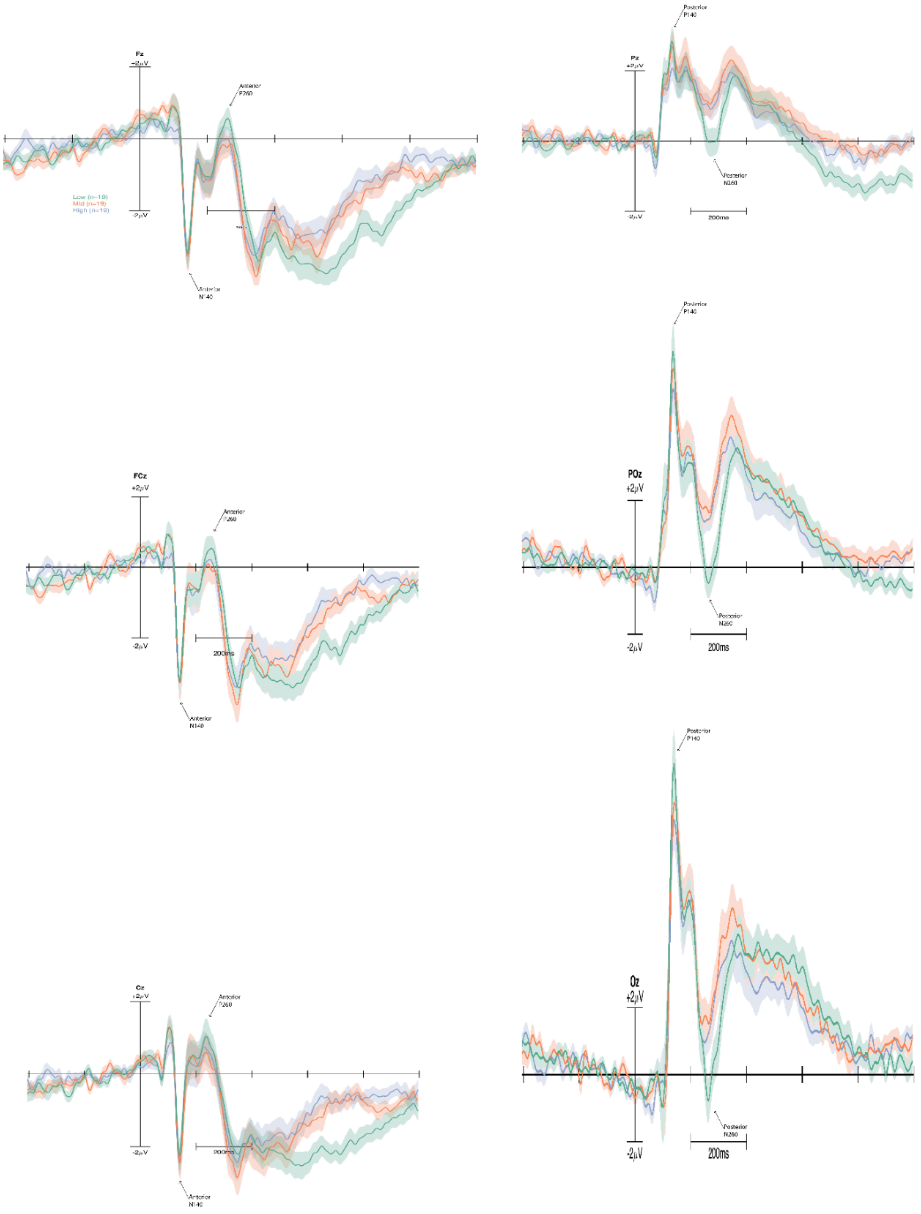


**Fig. S1.** Grand average event-related potentials with onset of the room display at all 6 electrodes. The different door heights are colour coded with the low door condition in green, the mid door condition in red and the high door condition in purple. Prominent components (Anterior N140, P260, Posterior P140, N260) as observed over electrodes Fz, FCz, Cz, Pz, POz and Oz are indicated by arrows.

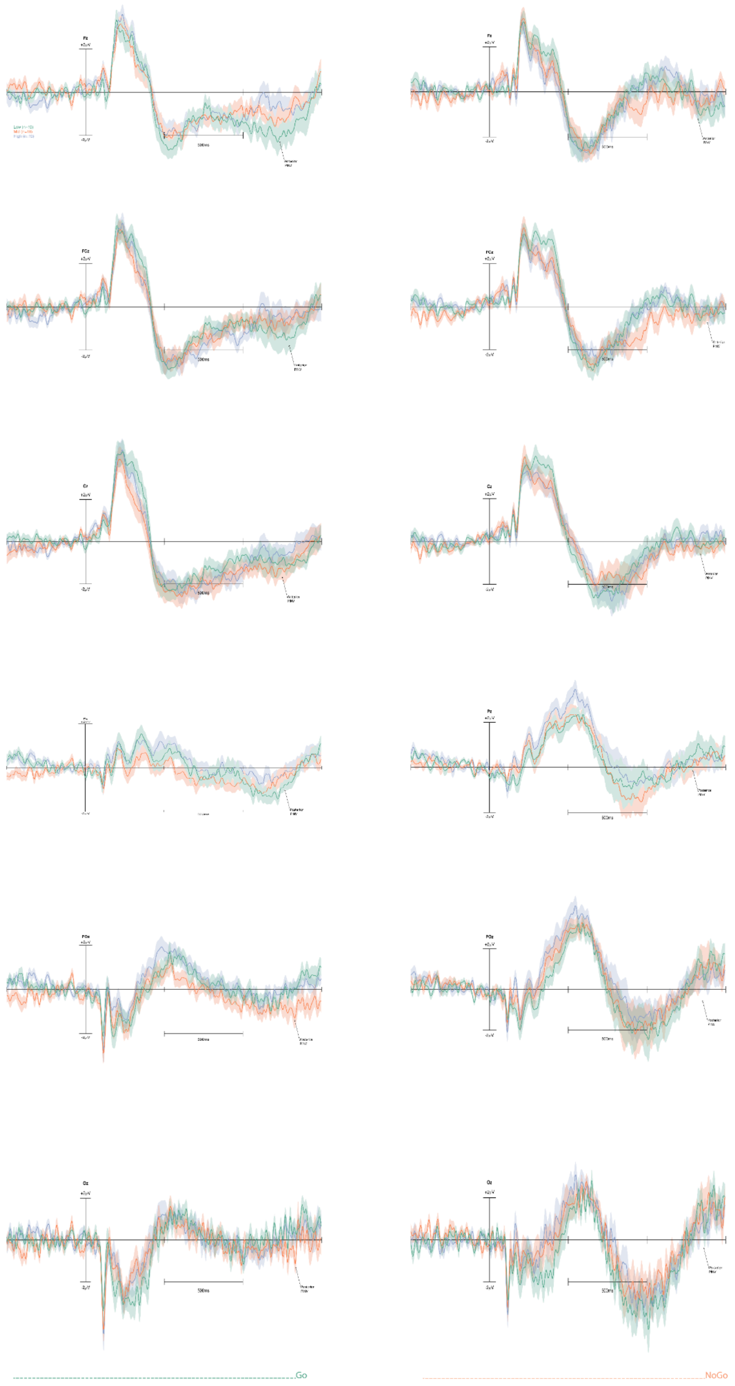


**Fig. S2.** Twelve time-locked event-related potentials (ERPs) from electrodes Fz, FCz, Cz, Pz, POz and Oz at the onset of imperative stimulus (Go, NoGo). The low condition is in green, the mid condition is in red, and the high condition is in purple. The plots on the left column showed the ERPs under the Go condition, whereas the plots on the right column showed the ERPs under the NoGo condition. PINV components were marked with arrows.

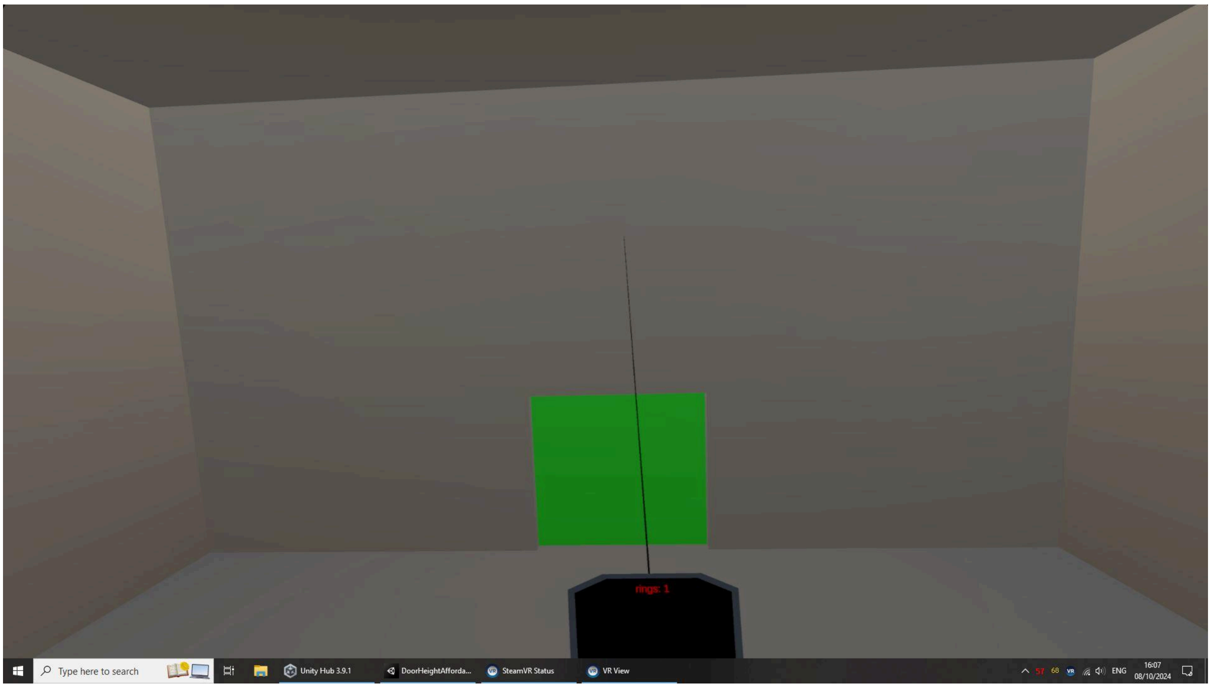


**Fig. S3.** Low door green (Go trial)

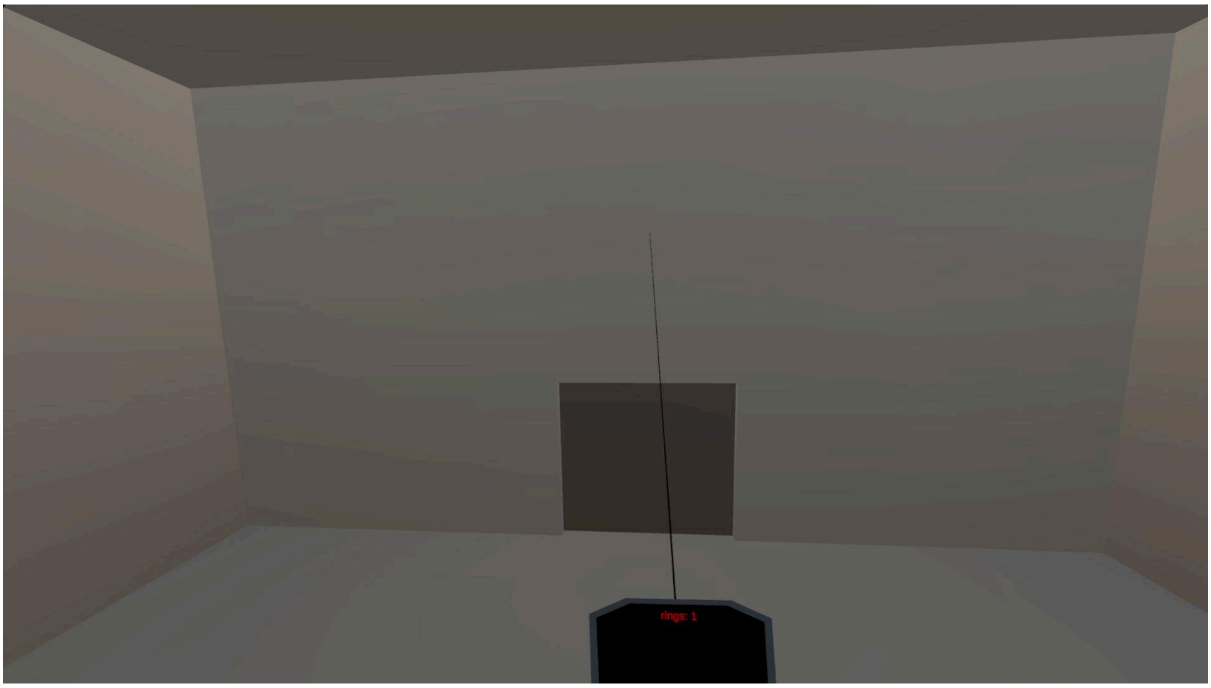


**Fig. S4.** Low door grey (lightsOn)

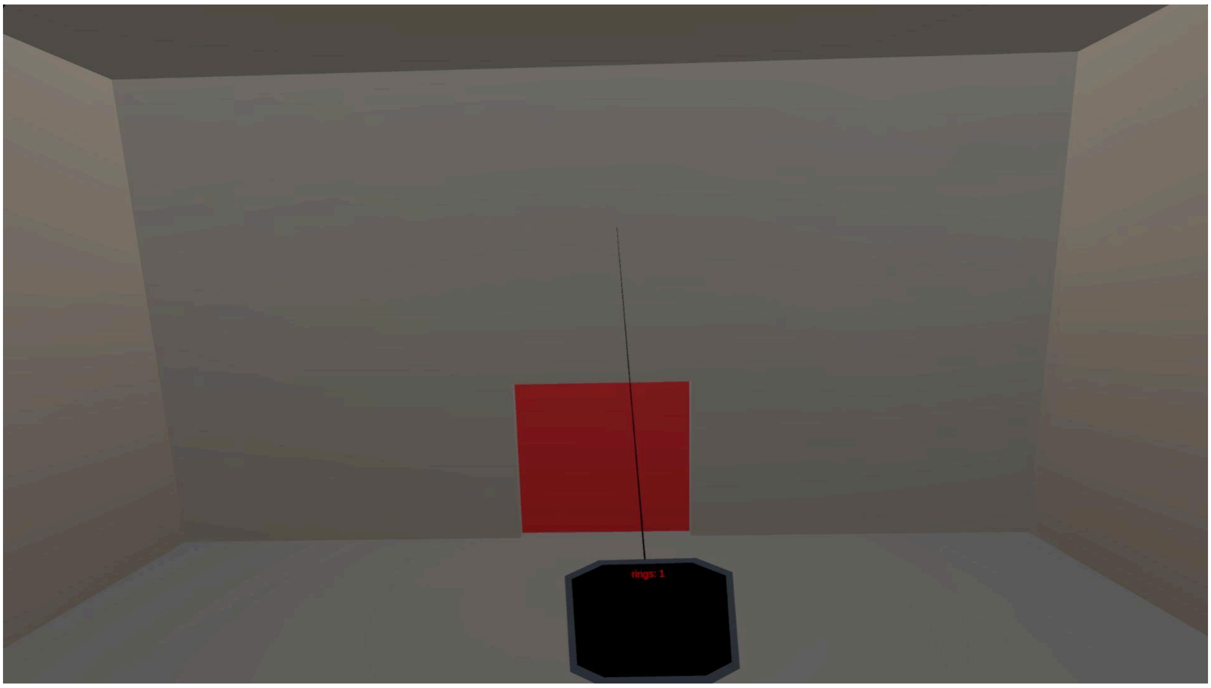


**Fig. S5.** Low door red (NoGo trial)

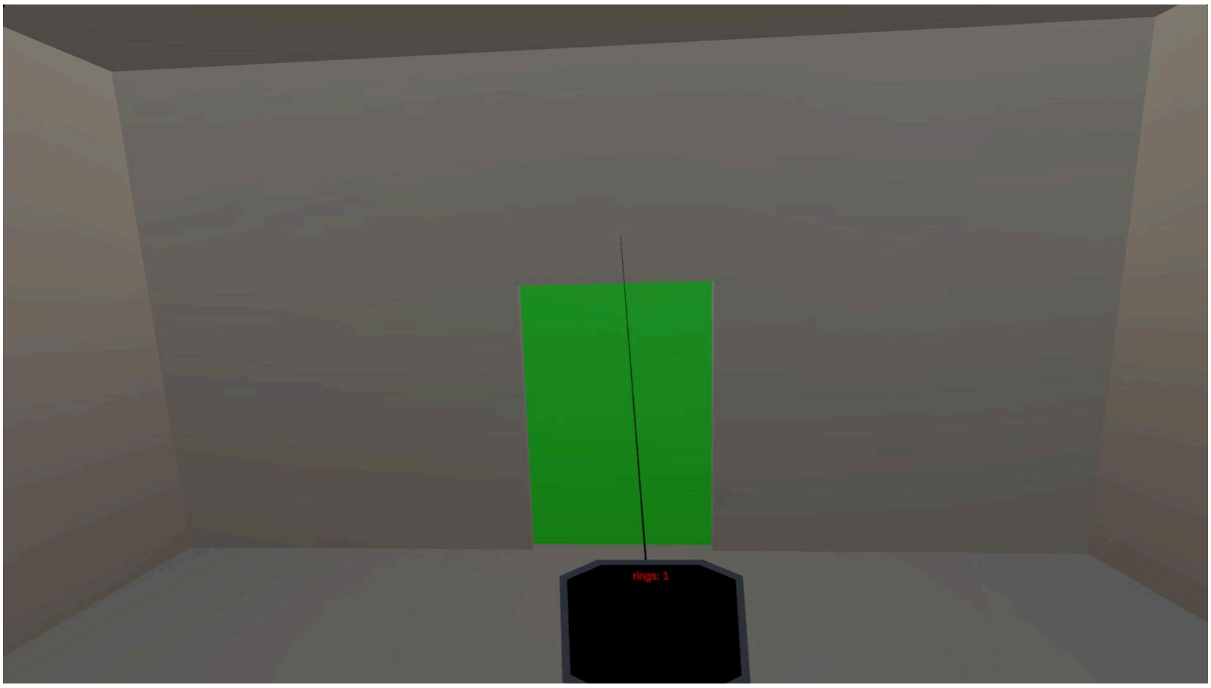


**Fig. S6.** Mid door green (Go trial)

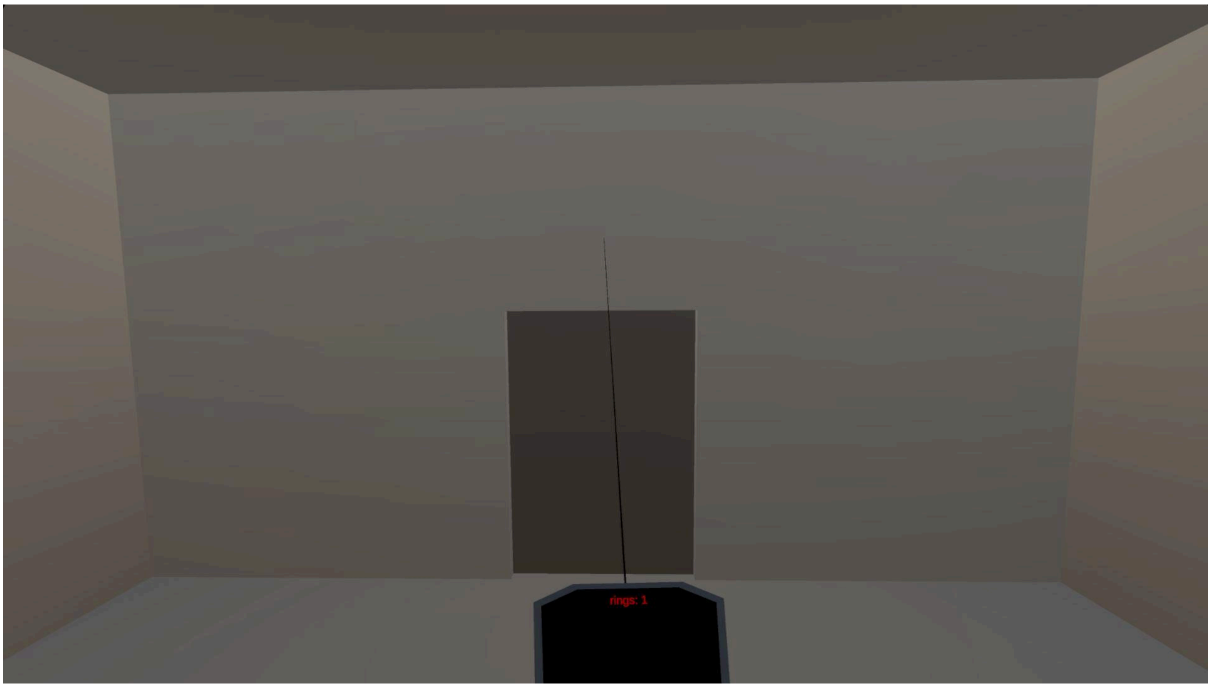


**Fig. S7.** Mid door grey (lightsOn)

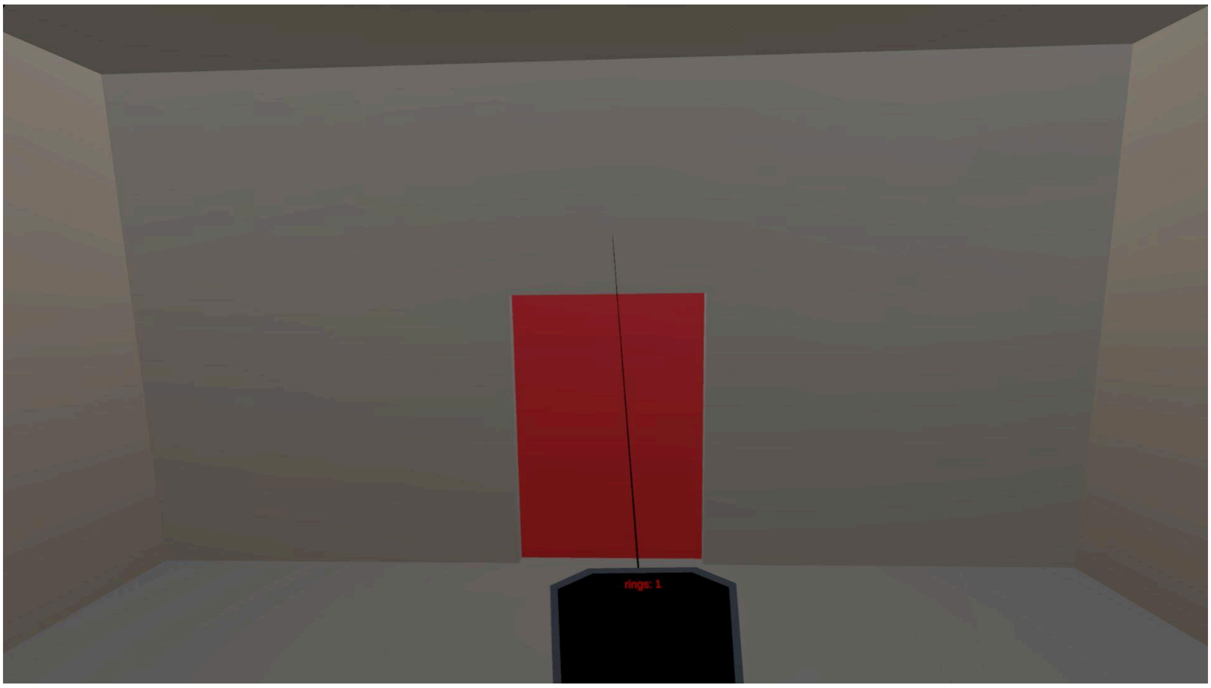


**Fig. S8.** Mid door red (NoGo trial)

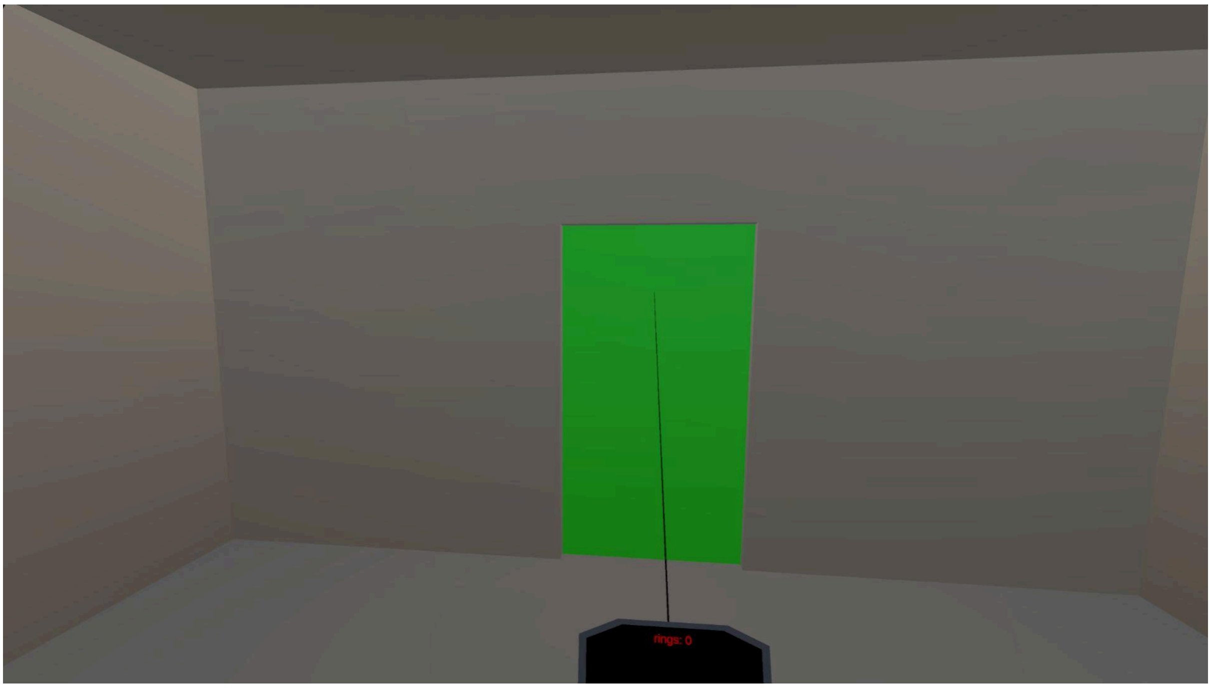


**Fig. S9.** High door green (Go trial)

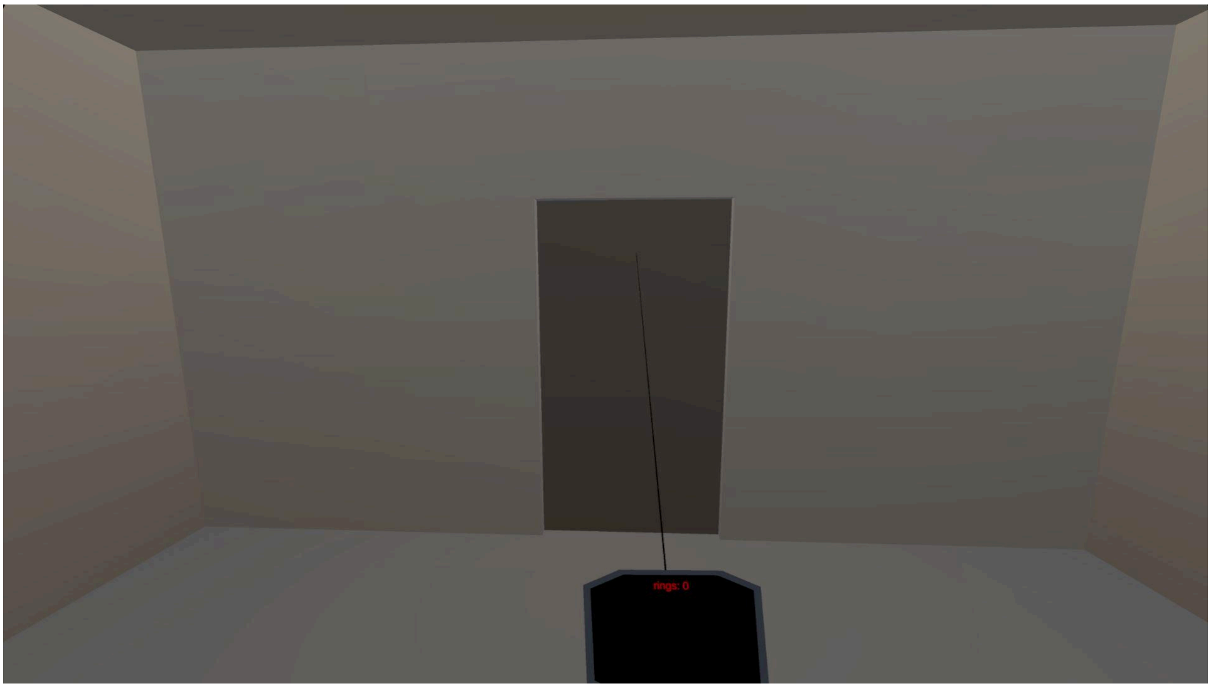


**Fig. S10.** High door grey (lightsOn)

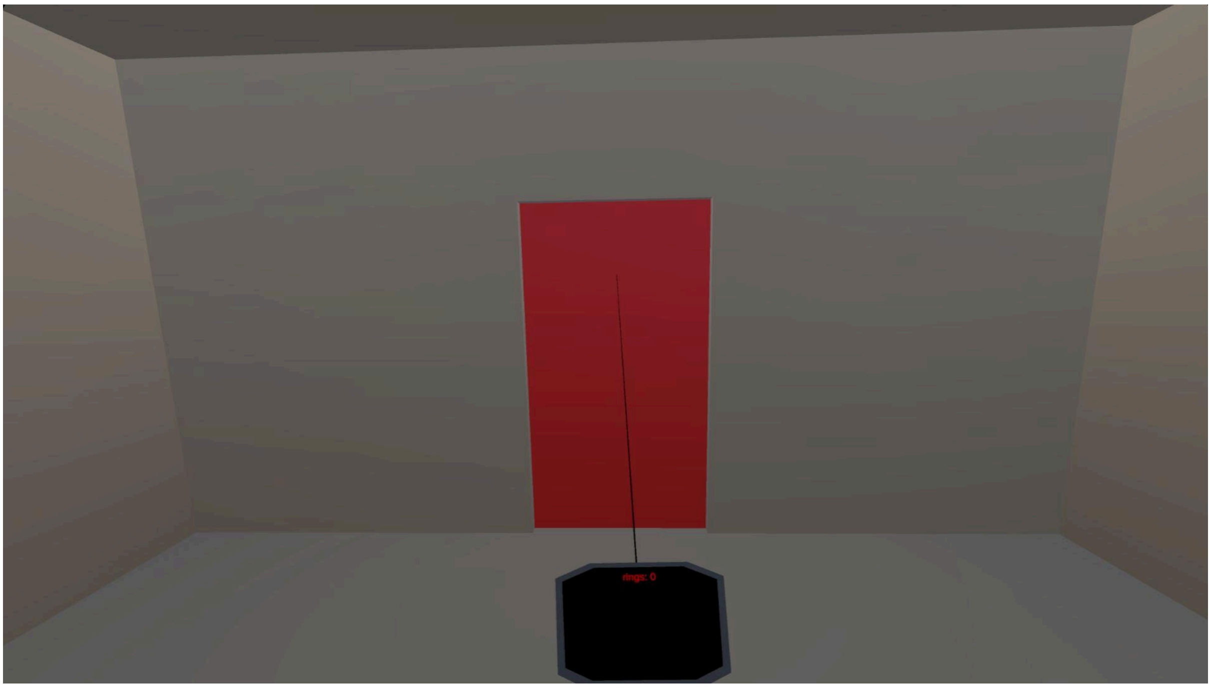

**Fig. S11.** High door red (NoGo trial)

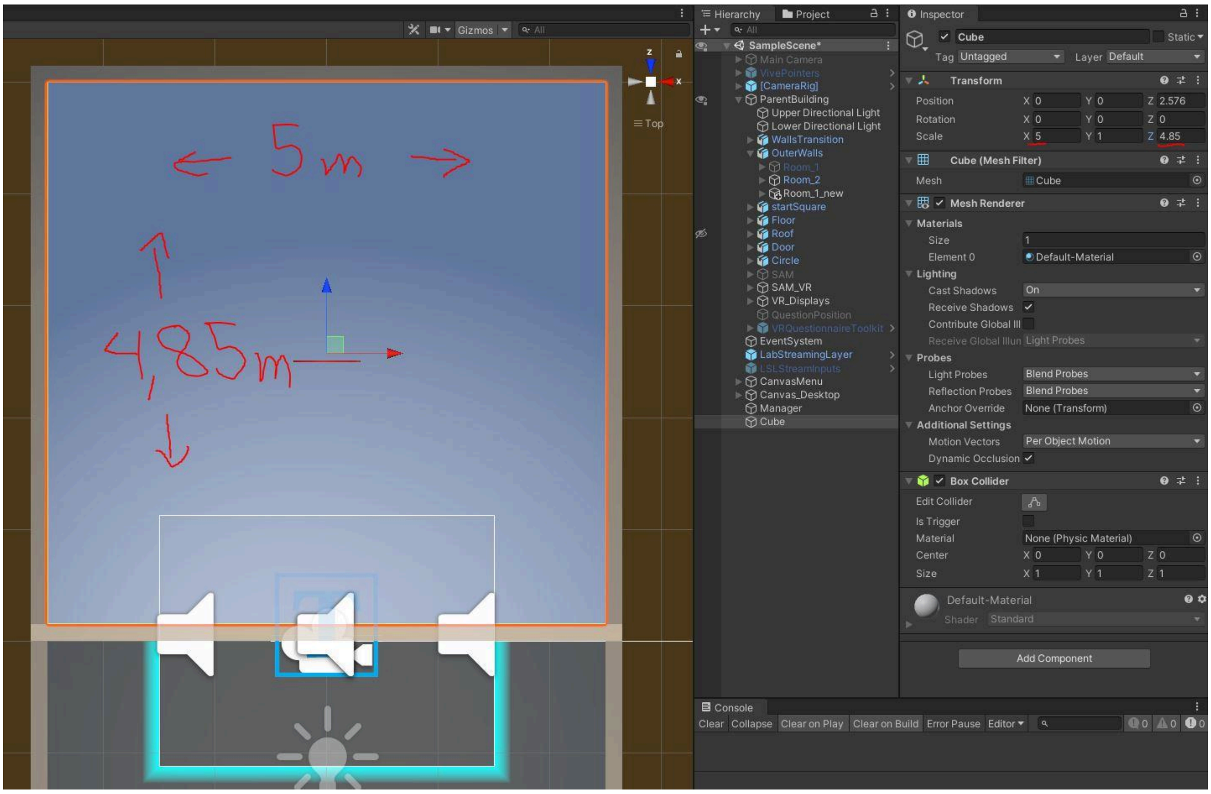


**Fig. S12.** Second room (Unity layout)

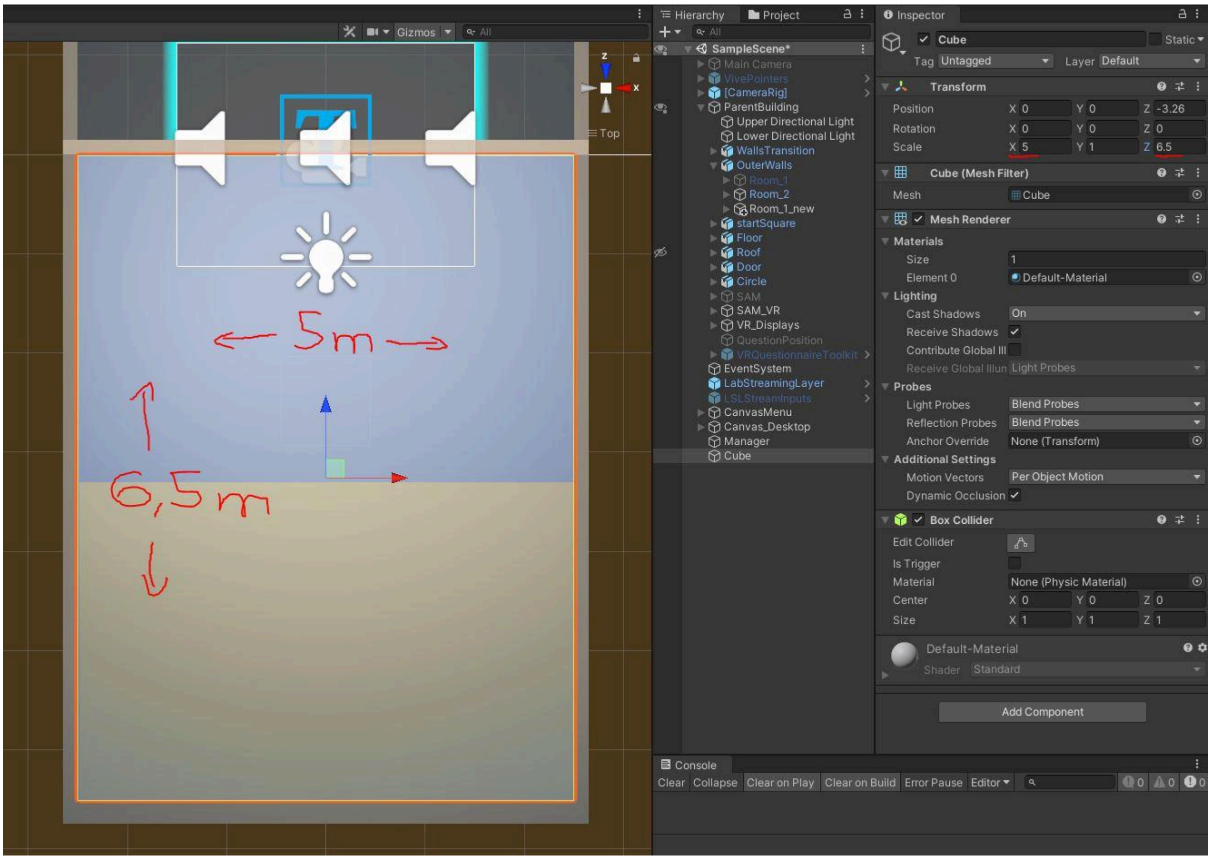


**Fig. S13.** Starting room (Unity layout)
